# Contrasting Modes of Dosage Compensation on Ancestral and Neo-Z Chromosomes in Butterflies

**DOI:** 10.64898/2026.07.30.741693

**Authors:** Lars Höök, Jesper Boman, Karin Näsvall, Daria Shipilina, Roger Vila, Niclas Backström

## Abstract

The evolution of sex chromosomes from homologous autosomes generally causes degeneration of the sex-limited chromosome (Y in XY systems and W in ZW systems), leading to altered or disrupted gene regulation. Dosage compensation has evolved repeatedly to mitigate the effects of such unbalanced expression of sex-linked genes. Previous analyses show that X chromosomes often are upregulated in males in XY-systems, while upregulation of the Z chromosome in females is unusual in ZW-systems, but understanding the generality of this pattern requires broader taxonomic sampling. In addition, little is known about how dosage compensation is established when sex chromosomes initially evolve. Here, we narrow this knowledge gap by characterizing dosage compensation in butterflies (*Leptidea* sp.) that harbor both ancestral, intermediate and recently derived sex chromosomes. Analyses of 120 samples, representing both sexes, different tissues and developmental stages, reveal a mixture of gene expression patterns. We confirm that downregulation in males is the predominant mode on the ancestral Z chromosome, but we find complex patterns of male downregulation and female upregulation for Z chromosomes with different evolutionary histories, as well as between different tissues. Dosage compensation has for example evolved rapidly in regions of the youngest neo-Z where the neo-W gametologs have lost function, but not in regions with functional gene copies on both Z and W. Our results provide novel insights into the complex evolutionary trajectories underlying dosage compensation and show that development of specific mechanisms likely depends on both sex chromosome system and dosage sensitivity of sex chromosome-linked genes.

## 1 INTRODUCTION

In species with genetic sex determination, the development of a specific sex is usually initiated early in embryogenesis and depends on the presence of dimorphic sex chromosomes (Mittwoch, 1996). Sex chromosomes originate from ancestral homologous autosome pairs that become differentiated through sex-specific selection, resulting in sex chromosome systems where one sex is homogametic (ZZ males or XX females) and the other is heterogametic (ZW females or XY males) (Wright et al., 2016). Sex chromosome dimorphism is hypothesized to be the result of selection for reduced recombination to maintain linkage between a sex determining locus and sexually antagonistic alleles, which over time causes the sex chromosomes to differentiate (Ellegren, 2011). Another consequence of the strong linkage is a reduced efficacy to remove deleterious mutations due to selective interference (Otto, 2021). Genes that are not under strong purifying selection are therefore expected to degenerate on the sex-specific chromosome over time (Charlesworth, 2021). In addition, selective interference can also cause degeneration of regulatory sequences, resulting in a gradual reduction in gene expression levels, which in turn leads to reduced exposure to selection and, consequently, accumulation of deleterious mutations in coding sequences (Lenormand et al., 2020). Conversely, highly expressed genes are expected, and observed, to be more constrained and have a higher resistance against degeneration on the sex chromosomes (Kaiser et al., 2011; Charlesworth, 2021). The degeneration of the sex-specific chromosome will result in hemizygosity of many (or all) sex-linked genes in the heterogametic sex. The loss of the gene copy on the sex-limited chromosome can reduce gene expression levels and disrupt gene interactions that were established before the sex chromosomes started to differentiate (Graves, 2016). It is therefore expected that dosage compensation mechanisms should evolve to restore ancestral expression levels of sex-linked genes (Charlesworth, 1996). Early studies of dosage compensation in *Drosophila* showed that mechanisms have evolved to specifically modulate the activity of sex-linked genes by upregulation in the heterogametic males, resulting in a general balance between sex-linked and autosomal genes (Mukherjee and Beermann, 1965; Xie et al., 2025). Consequently, the classical model of dosage compensation evolution stressed the importance of restoring the gene expression levels between the sex chromosomes and the autosomes (Gartler, 2014). However, increased taxonomic sampling and more detailed gene expression profiling has put this canonical model into question (Mank, 2013). In particular, maintenance of a dosage compensation between sex-linked and autosomal genes may not always be critical, since this appears to be a less common situation than previously thought (Gu and Walters, 2017). In addition, no chromosome-wide regulation appears to occur in the male heterogametic platypus (Julien et al., 2012) or in the female heterogametic chicken, where only subsets of genes are dosage compensated (Ellegren et al., 2007). Sex chromosome gene expression can also vary between different developmental stages (Nozawa et al., 2014), tissues (Mank and Ellegren, 2009), and closely related species (Huylmans et al., 2017), complicating interpretations of dosage compensation mechanisms. In addition, sex chromosomes are often enriched for genes with sex-specific functions (Mongue and Walters, 2018) and sex-biased expression (Höök et al., 2019), which could influence the evolution of dosage compensation, but also confound analyses based on chromosome-wide estimates of expression levels (Huylmans et al., 2017).

The observed diversity of dosage compensation mechanisms has spurred new questions about what evolutionary forces are important for causing different mechanisms to evolve. Complete dosage compensation, where expression is fully balanced between both sexes and between sex chromosomes and the autosomes, has for example more often been observed in male heterogametic systems (Mullon et al., 2015). In contrast, female heterogametic systems appear to more often have partial dosage compensation, where only subsets of dosage-sensitive genes get compensated (Mank and Ellegren, 2009). This difference could potentially be related to an expected faster degeneration of Y chromosomes compared to W chromosomes due to male-biased mutation rates (Naurin et al., 2010). The fast degeneration of the Y could limit the potential to evolve a gene-wise upregulation on the X in male heterogametic systems, and force chromosome-wide mechanisms to be established, such as the X chromosome inactivation (XCI) system in mammals (Naurin et al., 2010). An alternative hypothesis is that the generally reduced adaptive potential of Z compared to X chromosomes, due to higher variance in male reproductive success, could make chromosome-wide dosage compensation less likely to evolve in female heterogametic systems (Mullon et al., 2015). Z chromosome regulation would instead be restricted to individual genes that are under strong purifying selection for specific expression levels (Mank, 2013). However, disentangling the impact of individual evolutionary forces remains a challenge. Another challenge is that comparative approaches often rely on contrasts between species with deep divergence. In such cases, the selective regimes that were present when dosage compensation became established might differ from the current situation and could be difficult to infer. An alternative strategy for broadening the understanding of what drives specific dosage compensation mechanisms to evolve, is therefore to study species where neo-sex chromosomes have formed more recently through fusions between autosomes and sex chromosomes (Nozawa et al., 2018). This can also reveal if conserved lineage-specific traits, such as heterogamety, can predict how specific dosage compensation mechanisms evolve.

The order Lepidoptera (moths and butterflies) has received increased attention in studies about sex chromosome evolution and serves as a useful complement to other female heterogametic systems. In addition, the relatively frequently occurring fusions between autosomes and both the ancestral Z and W chromosomes make lepidopteran species useful for studying the establishment of dosage compensation on neo-sex chromosomes (Rueda-M et al., 2024; Wright et al., 2024; Yoshido et al., 2020; Kalita and Keller Valsecchi, 2024; Xie et al., 2025). Previous studies on dosage compensation on the ancestral Z chromosome in both moths and butterflies have revealed a general pattern, where males have reduced expression of Z-linked genes compared to the autosomal average, largely balanced with the expression of genes on the single Z chromosome in females (Catalán et al., 2018; Gu et al., 2017, 2019; Höök et al., 2019; Huylmans et al., 2017; Walters et al., 2015; Kalita and Keller Valsecchi, 2024; Xie et al., 2025) – a state that is sometimes referred to as dosage balance (Kalita and Keller Valsecchi, 2024). Much less is known about dosage compensation on neo-Z chromosomes but results so far suggest that unique mechanisms can evolve to regulate expression on these chromosomes (Gu et al., 2017, 2019; Kalita and Keller Valsecchi, 2024; Xie et al., 2025). An attractive system for studying the evolution of dosage compensation on neo-sex chromosomes is the cryptic wood white butterflies in the genus *Leptidea*. The clade, formed by the three species *L. juvernica, L. sinapis* and *L. reali*, has recently undergone an extensive genome restructuring, which has resulted in the formation of several neo-sex chromosomes (Šíchová et al., 2015). While the Z-linked gene content is shared between the species and mostly collinear, there have been additional fissions, fusions and translocations in *L. juvernica* and *L. reali*, resulting in unique Z chromosome morphologies in each respective species (Hö ö k et al., 2023; Yoshido et al., 2020; Thö rn et al., 2026). The recruitment of autosomal fragments to the Z chromosomes appears to have occurred in a stepwise manner before and during the radiation of *Leptidea* and the W chromosomes consequently show different levels of degeneration (Yoshido et al., 2020; Höök et al., 2024). This means that the different Z chromosome regions represent distinct snapshots where the temporal dynamics of dosage compensation evolution can be characterized.

In this study, we investigate how gene regulation has evolved on butterfly sex chromosomes, with a particular focus on the neo-Z chromosomes in the *Leptidea* clade. It has previously been shown that dosage compensation on the ancestral Z in *L. sinapis* occurs through downregulation in males to match the expression level in females (Höök et al., 2019). Here, we analyze if similar forms of dosage compensation have evolved on the neo-Z chromosomes, and if the mechanisms are conserved in the closely related species *L. reali* and *L. juvernica*. Furthermore, previous results have indicated considerable sequence homology between genes on the most recently recruited neo-Z chromosome and corresponding neo-W gametologs (Hö ö k et al., 2023; Yoshido et al., 2020). This is a rare situation in Lepidoptera due to female achiasmy (lack of meiotic recombination), which should cause rapid degeneration of W-linked gametologs (Turner and Sheppard, 1975). We therefore investigate if putative W gametologs are expressed and how this affects the evolution of dosage compensation on the youngest Z chromosome.

## 2 METHODS

### RNA extraction and sequencing

RNA-sequencing data was generated for the three species *L. juvernica, L. reali* and *L. sinapis*. Mated female butterflies were caught in the field in Sweden (*L. juvernica* and *L. sinapis*) and Catalonia (*L. reali*) and kept in the lab for egg laying. F1 male and female offspring were sampled as instar V larvae (the day after the fourth molt) or as adults (the day after eclosion), and snap-frozen in liquid nitrogen. RNA was extracted separately from the head and the terminal five segments of the abdomen (containing the gonads) for each sample using Qiagen RNeasy Mini kits following standard protocols including DNase treatment. Five libraries of each sample group (3 *species*\*2 *sexes*\*2 *stages*\*2 *tissues = 120 libaries in total*) were prepared with a TruSeq stranded mRNA protocol and sequenced with Illumina NovaSeq6000 technology (150 bp paired-end reads) on one S4 lane by the National Genomics Institute (NGI) in Stockholm. The sequencing yielded a mean read count of 25.5 M per sample.

### Read trimming

Raw RNA-seq reads were quality trimmed with TrimGalore v.0.6.1 (Krueger, 2019) and Cutadapt v.4.0 (Martin, 2011). Poly A/T/N-tails and low complexity regions were trimmed using Prinseq v.0.20.4 (Schmieder and Edwards, 2011). Trimmed reads were filtered for contaminants using FastQ Screen v.0.15.2 (Wingett and Andrews, 2018). Unpaired reads remaining after filtering were removed using BBmap v.38.61b (Bushnell, 2019).

### Transcriptome assembly

In order to improve gene annotation, we generated *de novo* transcriptome assemblies with Trinity v.2.13.2 (Grabherr et al., 2011), a separate transcriptome was generated for each species, sex and developmental stage. Transcriptome qualities were assessed with two complementary methods. First, reads were mapped back to each respective transcriptome using bowtie2 v.2.5.1 (Langmead and Salzberg, 2012), which resulted in a high mapping rate for all samples (94.97 - 98.96%). Next, core gene completeness was assessed with BUSCO v.5.3.1 (Manni et al., 2021) using the lepidoptera odb10 lineage set, which resulted in high levels of completeness (90.6 - 98.1%) (Supplementary table 1). In addition, expression quantification was performed with Salmon v.1.10.1 (Patro et al., 2017) and Kallisto v.0.48.0 (Bray et al., 2016) and transcripts without detectable expression count (*<* 1) in any sample were filtered out. Finally, highly similar transcripts (*>* 99%) were deduplicated with cd-hit v.4.8.1 (Fu et al., 2012).

### Gene annotation and orthology prediction

Gene annotation was performed separately for the three available *Leptidea* genome assemblies (Höök et al., 2023) using the Maker v.3.01.04 pipeline (Cantarel et al., 2008). Maker was first run to map the generated transcripts and previously available protein sequences of each respective species (Höök et al., 2023), in order to generate initial gene models. Repeat masking was performed as part of the initial mapping step with Maker, using previously generated repeat libraries for each species (Höök et al., 2023). The gene models generated from the first run were then used to train the gene predictor Augustus v.3.4.0 (Stanke et al., 2008) and the predicted genes were incorporated in the second run of Maker. Gene prediction was then reiterated three more times to improve the gene prediction parameters. The final annotation resulted in the expected number of annotated genes based on previous annotations (Supplementary table 2) and generally low alignment edit distance (AED), suggesting high agreement between annotation predictions and evidence (Supplementary figure 1). Functional prediction was performed with Interproscan v.5.30-69.0 (Jones et al., 2014) and homology searches against the Swiss-Prot database (Bairoch and Apweiler, 2000) using blastp v.2.13.0 (Altschul et al., 1990). This resulted in functional annotations for *>* 9,509 genes in each species (annotation statistics are summarized in Supplementary table 2). Single-copy orthologs were identified using Orthofinder v.2.5.4 (Emms and Kelly, 2019) based on the longest isoform of each gene.

### Read mapping and differential gene expression analysis

RNA-seq reads were mapped to the male reference assembly of each respective species using Star v.2.7.9a (Dobin et al., 2013) in two-pass mode, guided by gene annotation. Read counts for each sample were quantified with the ‘featureCounts’ function in Subread v.2.0.3 (Liao et al., 2014). Sex-biased gene expression was analyzed separately for each species, stage and tissue using edgeR v.3.40.2 (Robinson et al., 2010). Genes with low expression levels were filtered using the ‘filterByExpr’ function and library scaling was performed with the ‘trimmed mean of m’ method (TMM). Differential expression was tested using the ‘glmTreat’ function with a cutoff level of |1.5 |on unshrunk log2 fold change values to detect differentially expressed genes, and a significance threshold for FDR adjusted p-value of *<* 0.05.

### Dosage compensation analysis

In *Leptidea*, different autosomal regions have become Z-linked in a stepwise manner (Höök et al., 2024). To analyze if different forms of dosage compensation mechanisms have evolved in the different neo-Z chromosome regions, we compared average expression levels of genes in each respective region to the expression levels of autosomal genes. Read counts of expression filtered and TMM-scaled libraries (see differential gene expression analysis) were normalized with the ‘transcripts per million’ method (TPM) to account for differences in length between gene sets. Genes without expression in a sample group were filtered out. Gene-wise mean TPM was calculated for each sample group and differences between sets of genes were evaluated with Kruskal-Wallis rank sum tests followed by pairwise Wilcoxon rank sum tests corrected for multiple testing using the Benjamini-Yekutieli method, to account for the statistical dependence between tests. In addition, Z/A ratios of median gene expression levels were calculated for each separate Z chromosome region, and confidence intervals of the ratios were calculated using stratified bootstrap, accounting for the difference in sample sizes between gene sets, using 10,000 replicates.

### Variant calling

Identification of SNPs was performed for the RNA-seq data for each species separately. Base quality score recalibration and joint variant calling of all sample groups were made with HaplotypeCaller in GATK v.4.1.1.0 (McKenna et al., 2010), following recommendations for RNAseq short variant discovery, and the resulting vcf files were filtered with bcftools v.1.14 (Li, 2011) using the filtering criteria ‘-I QUAL*>* 30 && QD*>* 2 && FS*<* 60 && MQ*>* 40 && FMT/DP*>* 5’. In addition, to make contrasts between observed (DNA reads) and expressed SNPs (RNA reads), variant calling of a previously available DNA resequencing data set with 10 male samples for each of the focal *Leptidea* species (Talla et al., 2019) was performed with HaplotypeCaller as described in (Höök et al., 2024).

### Detection and expression quantification of W gametologs

Expression of W-linked gametologs was analyzed by first identifying female-unique exonic SNPs in the RNA-seq data. The SNPs were identified using the following criteria: 1) the position was heterozygous in all female samples, 2) the position was homozygous for the reference allele in all male samples, 3) the position did not overlap repeat sequences in the male reference assembly, and 4) the position was not called as a variable site in a DNA resequencing data set which contain 10 male samples of each species (Talla et al., 2019). The analysis was performed by first using the whole set of female samples (10 *individuals*\*2 *tissues* of each sex per species, in total) resulting in a “core set” of female-unique SNPs, and then for each separate female sample group (*tissue*\**stage*), compared against all male samples. To assess the false discovery rate (i.e. the chance to randomly find unique SNPs in one sex) we identified male-unique SNPs in the RNA-seq data using the same criteria as described for detecting female-unique SNPs. All overlap assessments were performed with bedtools v.2.29.2 (Quinlan and Hall, 2010). Expression of Z-linked and W-linked alleles (i.e. relative allele-specific expression) was then estimated with phaser v.20210423-5d4926d (Castel et al., 2016) by comparing the reference allele (Z) and the female unique alternative allele (W) expression counts after filtering out genes, with *<* 10 counts for any allele to reduce false positives due to random, incorrect mappings. No normalization was performed for the estimated counts since they represent contrasting expression levels of alleles within individual libraries.

### GO analysis

A GO analysis was used to assess if certain functional categories were overrepresented among gene sets located in the different Z chromosome regions, using all Z-linked genes as the background set. Overrepresentation (p-value threshold *<* 0.05) of GO terms was estimated with TopGO v.2.50.0 (Alexa and Rahnenführer, 2023) using Fisher’s exact test and the ‘weight01’ pruning algorithm which combines the ‘elim’ and ‘weight’ algorithms to account for the GO topology.

## 3RESULTS

We analyzed 120 mRNA libraries of the three closely related species *L. sinapis, L. juvernica* and *L. reali* and quantified sex-linked and autosomal gene expression. Z-linked genes were partitioned into five different ‘Z regions’ (Z1a, Z1b, Z2a, Z2b, Z3) based on the inferred evolutionary history of the step-wise recruitment of neo-Z fragments (Figure 1; Höök et al., 2024). To compare equivalent gene sets across species, genes were grouped according to the Z chromosome karyotype of *L. sinapis* (Figure 1), i.e. the ancestral state in the clade. Note however that additional rearrangements have occurred in *L. juvernica* and *L. reali* (Supplementary figure 2). First, we categorized genes with sex-biased expression to assess the general dosage balance between the sexes and identify potential differences in sex-specific regulation between developmental stages and tissues. Such patterns are important to consider when interpreting patterns of dosage compensation. A clustering analysis in each respective species showed that abdomen samples always clustered by sex, while head samples only clustered by sex in the adults (Supplementary figure 3). However, the separation of abdomen samples was much more distinct, as expected since abdomens contain gonads (Supplementary figure 3). A differential gene expression analysis showed that very few genes had sex-biased expression in larval and adult heads (Supplementary table 3), and all genes were therefore included when comparing sex-linked and autosomal expression in these sample groups. The number of sex-biased genes was considerably higher in adult abdomen, with a higher proportion of male-biased than female-biased genes in both stages (Binomial tests, p *<* 2.20*10^-16^, Supplementary figure 4, Supplementary table 3). Due to this difference in proportions of sex-biased genes, dosage compensation was assessed both including and excluding the sex-biased genes in adult abdomen.

**Figure 1.**
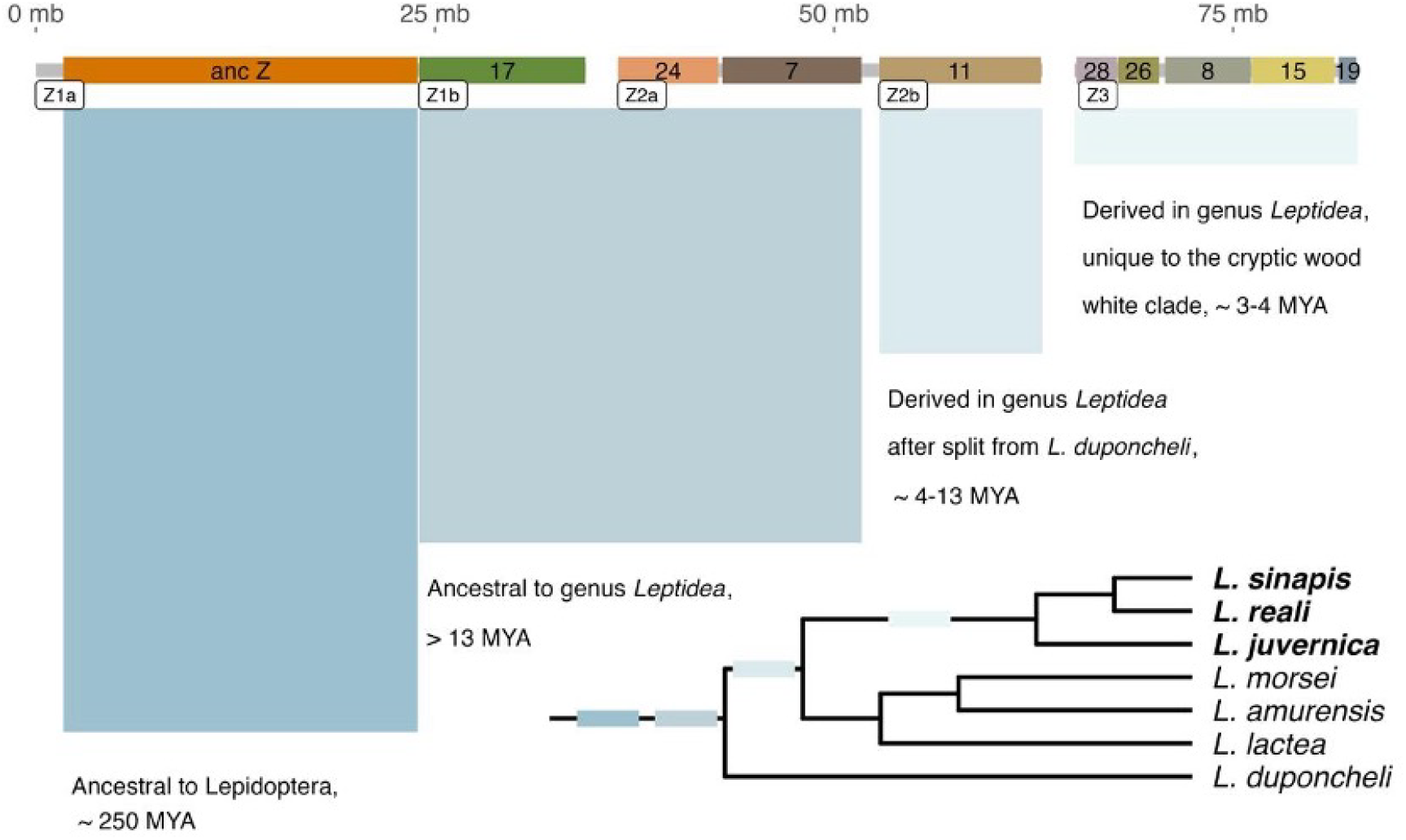
Inferred evolutionary history of neo-Z chromosomes in *Leptidea*. The chromosomes represent the ancestral Z chromosome karyotype of the genus, which is conserved in *L. sinapis*. Z chromosome fragments are colored / labelled based on homologous chromosome regions in *B. mori* (grey regions are not part of inferred synteny blocks). The white labels under each chromosome show the five Z chromosome regions that were analyzed separately in this study (Z1a, Z1b, Z2a, Z2b, Z3). The cladogram shows the *Leptidea* clade with the focal species highlighted in bold, and shaded boxes indicate along which branch the different Z chromosomes have formed. The topology of the cladogram is based on (Dincă et al., 2011)

### Contrasting modes of dosage compensation on different Z chromosomes

We compared gene expression levels between the autosomes and the different Z chromosome regions and found significant differences in all sample group comparisons (Kruskal-Wallis tests, p *<* 2.070*10^-5^, Supplementary table 4). The analysis of genes on the ancestral Z chromosome region confirmed the previously described male downregulation in *L. sinapis* (Wilcoxon tests, p *<* 2.66*10^-5^), and this pattern was consistent across all sample groups also in *L. juvernica* (Wilcoxon tests, p *<* 0.002) and *L. reali* (Wilcoxon tests, p *<* 0.013; Figure 2, Supplementary figure 5, Supplementary tables 5-10). In contrast, genes on the neo-Z region fused to the ancestral Z (Z1b, homologous to part of *B. mori* autosome 17) did not have significantly different expression levels compared to the autosomes in either sex in any of the species or sample groups (Wilcoxon tests, p *>* 0.088; Figure 2, Supplementary figure 5, Supplementary tables 5-10), showing that genes located on Z1b are upregulated in females, i.e. that complete dosage compensation has evolved for this Z region.

**Figure 2.**
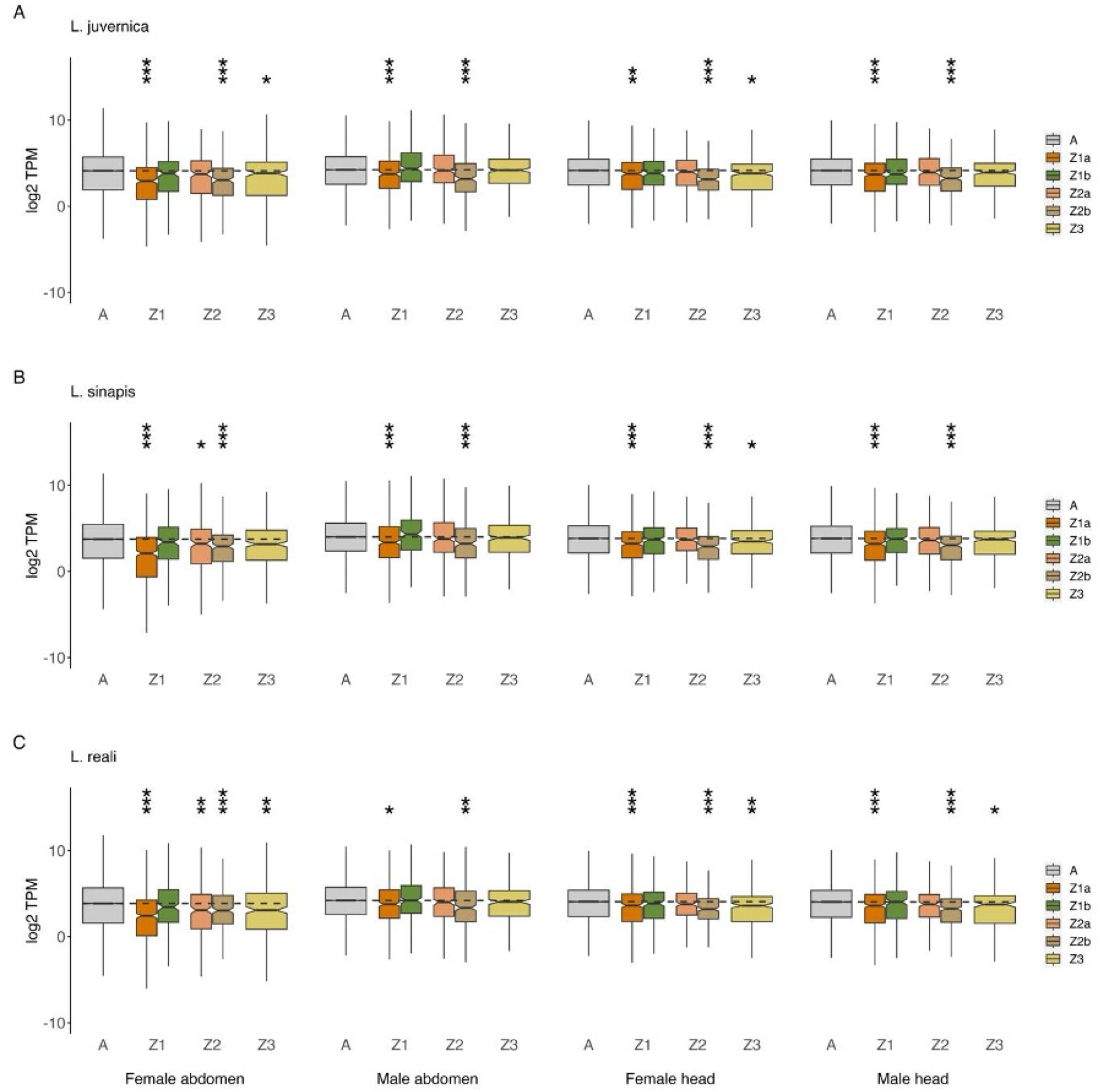
Comparisons of expression levels in adults between the five different Z chromosome regions and the autosomes. Note the strong concordance in results among species and tissues. The black dashed lines show median autosomal levels for each sample group. Asterisks indicate significantly reduced expression levels for Z regions compared to the autosomes (* P ⩽ 0.05; *∗∗ P* ⩽ 0.01; *∗∗∗ P* ⩽ 0.001. P-values and test statistics for all pairwise comparisons are available in Supplementary tables 6-8 and Z / A median expression ratios are available in Supplementary tables 10-12.

The genes on the second Z chromosome (Z2, which has shared synteny between all three *Leptidea* species and is homologous to parts of *B. mori* chromosomes 24, 7 and 11) had a mixed pattern in larvae and a reduced expression compared to autosomal genes in general (Supplementary figure 5, Supplementary tables 5-10). The genes on the more recently recruited region on this chromosome (Z2b, *B. mori* chromosome 11) had significantly reduced expression compared to autosomes in all sample groups (Wilcoxon tests, p *<* 0.002). The results for the older region of Z2 (Z2a, *B. mori* chromosomes 7 and 24) were more variable, with significantly reduced expression levels in all *L. reali* sample groups (Wilcoxon tests, p *<* 0.04), in females in general (Wilcoxon tests, p *<* 0.012) and in male abdomen (Wilcoxon test, p = 0.026) in *L. juvernica*, but only in male abdomen in *L. sinapis* (Wilcoxon test, p = 0.045; Supplementary figure 5, Supplementary tables 5-10). In adults, Z2 more consistently showed a similar form of dual dosage compensation as observed for Z1 (Figure 2). The genes on the Z2b region again had consistently reduced expression levels, suggesting male downregulation (Wilcoxon tests, p *<* 0.006; Figure 2, Supplementary tables 5-10). Conversely, the Z2a region was not significantly different from the autosomes, except in *L. sinapis* and *L. reali* female abdomen where expression levels were significantly reduced (Wilcoxon tests, p *<* 0.032; Figure 2, Supplementary tables 5-10). Taken together, this suggests that Z2a becomes dosage compensated (i.e. expressed at autosomal levels) in adults, but less so in female abdomen where expression is generally reduced. Importantly, this difference in expression levels between Z2a and the autosomes for female abdomen disappears when genes with sex-biased expression are excluded, while the pattern on Z2b is not affected by this filtering (Supplementary figure 6, Supplementary table 11).

The genes on the most recently acquired Z chromosome (Z3) did not have significantly reduced expression levels compared to autosomal genes in larvae, except for in head of *L. reali* females (Wilcoxon test, p = 0.042; Supplementary figure 5, Supplementary tables 5-10). In adults, the patterns for Z3 were more complex, with subtle differences between the species (Figure 2). In *L. reali*, genes on Z3 had reduced expression compared to autosomes in all sample groups (Wilcoxon tests, p *<* 0.027) except male abdomen (Wilcoxon test, p = 0.946). In *L. juvernica*, expression of genes located on Z3 was reduced in females (Wilcoxon tests, p *<* 0.028), but not in males (Wilcoxon tests, p *>* 0.23), while in *L. sinapis*, female head was the only sample group with significantly reduced expression (Wilcoxon test, p = 0.026; Figure 2, Supplementary tables 5-10). Importantly, the average expression levels for genes on Z3 were generally higher than for genes on both the ancestral Z (Z1a) and the Z2b region (Figure 2, Supplementary tables 5-10). In addition, there was no significant difference in expression levels between Z3 and autosomal genes in adult abdomen after removing genes with sex-biased expression (Supplementary figure 6, Supplementary table 11). In summary, the genes located on Z3 were generally expressed at autosomal levels, but with subtle differences between the species and with a tendency for female downregulation in general.

### Evolution of dosage compensation and expression of W-linked gametologs

The most recently acquired neo-Z region (Z3) provides a prominent case to characterize patterns of gene expression at an early stage of sex-chromosome differentiation, since the homologous neo-W chromosome (W3) still harbors multiple gametologous genes. Our initial analysis revealed that most sample groups had a near complete balance in Z3 gene expression between males and females, and between Z3 genes and autosomal genes. We therefore wanted to investigate if this pattern was caused by upregulation of Z3-linked genes in females (i.e. dosage compensation) or if some W3 gametologs are expressed, resulting in dosage balance. First, we identified Z3-linked genes that had female-specific polymorphisms (i.e. SNPs shared by all female individuals and absent in males), which would indicate expression of a W3-linked gametolog. When combining all samples (head and abdomen samples from 10 individuals) for each sex per species, we found between 274 - 338 female-unique SNPs distributed across 59 - 61 genes that were almost exclusively located on Z3 (Supplementary table 12). It should be noted that no male-specific SNPs were found in any of the species when combining all tissues and stages, indicating that the risk to erroneously score sex-specific SNPs should be low in our sample set. When analyzing tissues and stages independently in female samples, we detected a set of genes with female-unique SNPs shared across all sample groups (Supplementary table 13). The majority of female-unique SNPs occurred in all stages and tissues, with a tendency for more unique genes in head tissue (Supplementary figure 7). When including data from all sample groups, we identified 56 single-copy orthologs with female-unique SNPs present across all three species, a significantly higher overlap compared to a random resampling distribution of overlapping Z3 orthologs (Resampling test, p = 1.0*10^-4^; Supplementary figure 8). These results provide evidence for expression of at least 56 W3 gametologs in females that are conserved between the *Leptidea* species.

To assess the level and breadth of expression of the identified W3 gametologs, we analyzed the allelic imbalance (Z3 / W3 expression ratio in females) as compared to the expression of homologous genes on Z3 in males. If the expression of Z3 gametologs compensates for reduced W3 expression, we expect to see a comparatively high expression of the Z3 compared to the W3 gametolog in females. We tested this prediction by assessing the correlation between the male / female gene expression ratio (log2 fold change) and the expression ratio between the Z3 and W3 alleles in females. Contrary to our prediction, we found a positive correlation between these ratios, showing that lower expression levels in females relative to males is associated with reduced expression of W3 gametologs (Figure 3). This correlation was significant for all species, tissues and developmental stages except in adult abdomen where there was no correlation in any species (Figure 3, Supplementary table 14). This indicates a lack of a dosage compensation for genes on Z3, or that female-specific regulation occurs for certain genes. We found no equivalent correlation between allele-specific expression in males and the male / female gene expression balance, except for a weak negative correlation in *L. juvernica* larval abdomen (Figure 3, Supplementary table 14). In addition, there was no significant correlation between absolute gene expression levels and allele-specific expression in females (Supplementary figure 9, Supplementary table 15).

**Figure 3.**
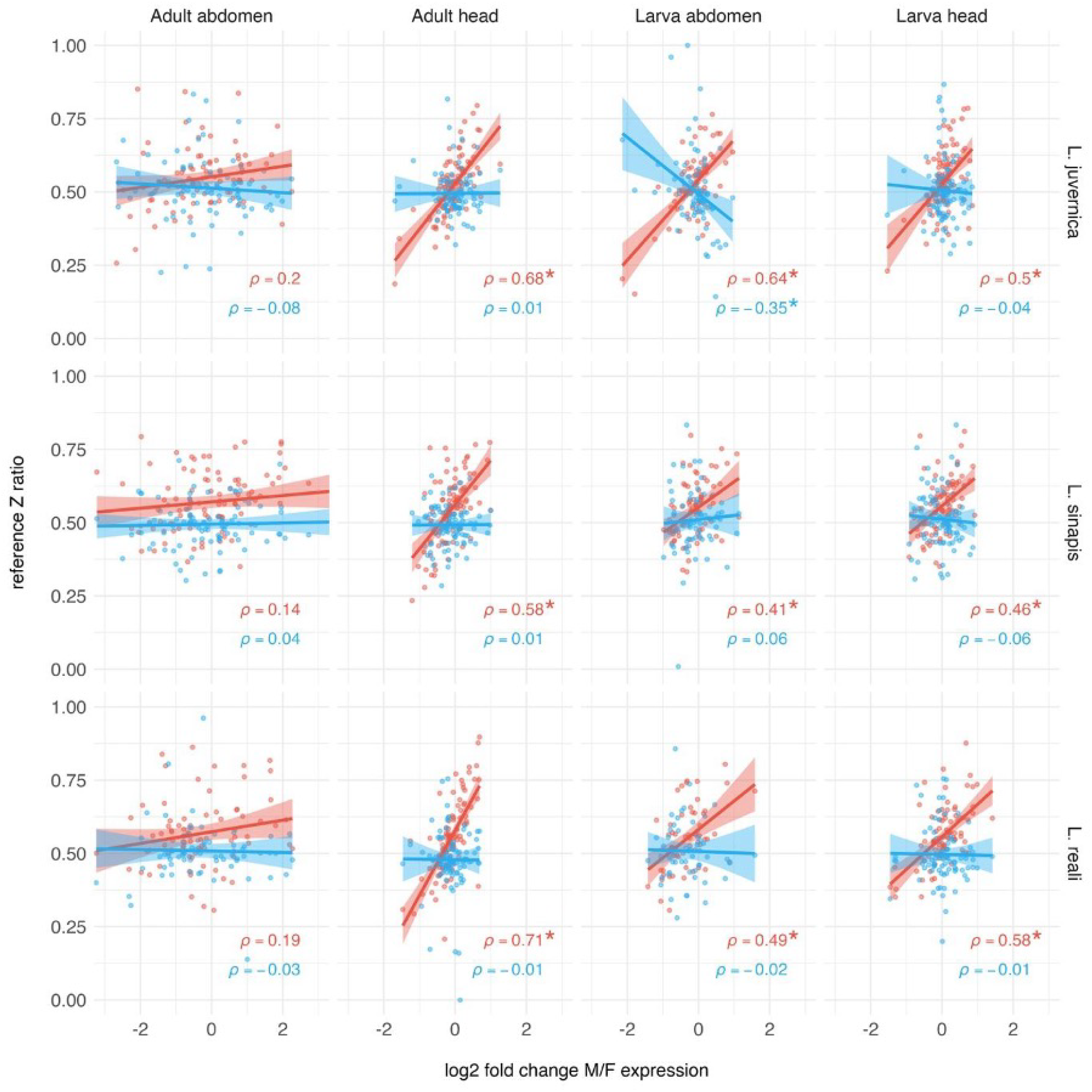
Associations between the male (M) / female (F) expression ratio (log2 fold change, X-axis) and the relative expression of the reference allele (Z / W in females, Z / Z in males, Y-axis) for gametologous genes on the Z3 and W3 chromosomes. The lines show the inferred regression for females (red) and males (blue). Significant correlations are indicated with asterisks. Test statistics and p-values are available in Supplementary table 14.

Next, we analyzed the expression levels for genes on Z3 that did not show any evidence for expression of W-linked gametologs. This subset makes up ∼ 46-71% (across different stages and tissues) of all genes on Z3 and could potentially have evolved dosage compensation in response to a lack of expression of the W-linked gametologs. Our previous observation of a mostly complete balance of expression levels between Z3 and the autosomes (Figure 2) suggests that upregulation has evolved in females for this set of genes. However, in contrast to those expectations, we found that genes where only the Z-linked allele is expressed in females (i.e. monoallelic expression; Z / 0) were dosage balanced between the sexes by a significantly reduced expression in males (compared to autosomal levels) in all sample groups (Wilcoxon tests, p *<* 0.014), except in *L. sinapis* larval abdomen (Wilcoxon test, p = 0.087; Figure 4A, Supplementary table 16). In addition, genes with bi-allelic expression (Z / W) were consistently expressed at a higher level than genes with monoallelic expression (Z / 0) in females in all comparisons (Wilcoxon tests, p *<* 3.390*10^-10^, Figure 4A, Supplementary table 17). We also found that the homologous Z3 genes in males (i.e. genes with bi-allelic expression in females) had higher expression than autosomal genes in males (Figure 4A, Supplementary table 17). Genes with bi-allelic expression in females (Z / W) were distributed across most of the Z3 chromosome and occurred interchangeably with genes showing monoallelic expression (Z / 0) (Figure 4B). These results suggest that dosage balance has evolved rapidly for the subset of genes where only the Z-linked allele is expressed in females, and that expression is regulated on a gene-by-gene basis, or in smaller clusters of genes, along this neo-Z region in males.

**Figure 4.**
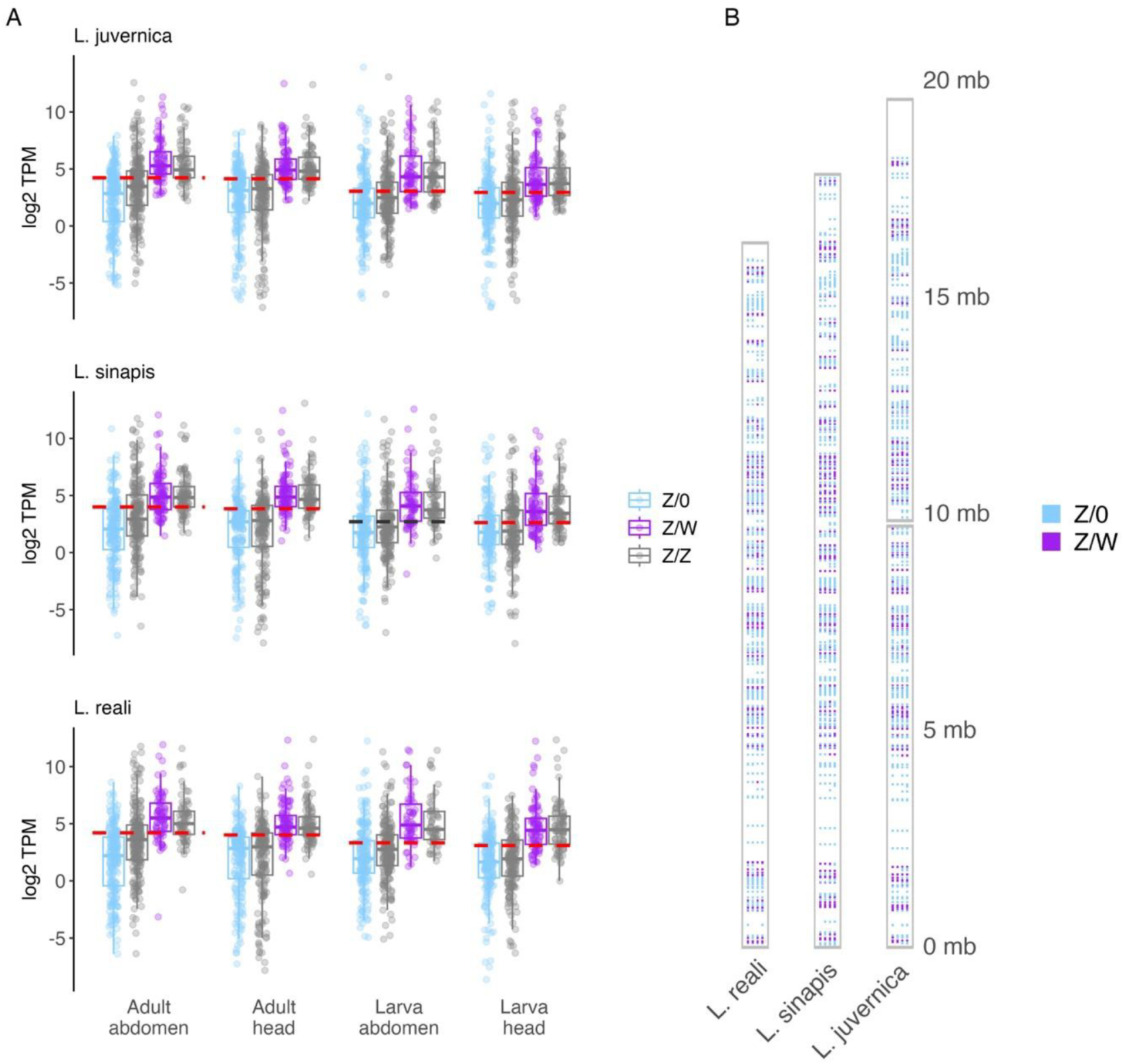
A. Distributions of expression levels for genes on chromosome Z3 with monoallelic expression (Z / 0) and bi-allelic expression (Z / W) in females. Genes with bi-allelic expression in females (purple) had significantly higher expression levels compared to genes with monoallelic expression (light blue) in all cases. The grey boxes show expression levels of the corresponding gene sets in males (Z / Z). The dashed lines show average expression of autosomal genes in males. Male expression levels of the bi-allelic gene set (i.e. the gene set where both gametologs were expressed in females) were also significantly higher compared to the autosomes in all cases (indicated with red dashed lines) except in *L. sinapis* larval abdomen (indicated with black dashed lines). B. Physical positions of genes with monoallelic (Z / 0) and bi-allelic (Z / W) expression on chromosome Z3. The sample groups are ordered from left to right: adult abdomen, adult head, larva abdomen, larva head.

### Gene ontology enrichment analysis

To understand if different modes of dosage compensation are associated with certain gene functions, we performed gene ontology (GO) analyses of Z-linked genes in regions showing either male downregulation or female upregulation. As expected, the results were largely overlapping between species, and we therefore interpret them collectively. Genes in downregulated Z regions (Z1a, Z2b and genes on Z3 with mono-allelic expression in females) showed overrepresentation of terms related to for example carbohydrate metabolism (GO:0005975 ‘carbohydrate metabolic process’) and transcription (GO:0003700 ‘DNA-binding transcription factor activity’) (Supplementary table 18). We also found enrichment of genes associated with ‘microtubule-based movement’ (GO:0007018), which is expected since spermatogenesis related genes are overrepresented on the ancestral Z chromosomes (Mongue & Walters, 2018). The most significant terms in Z regions with female upregulation (Z1b and Z2a) were related to sulfate transport (GO:0008272 ‘sulfate transport’, GO:0008271 ‘secondary active sulfate transmembrane transporter activity’) (Supplementary table 19). In addition, we analyzed enrichment of GO terms for genes on Z3 with expression of W-linked gametologs. This analysis resulted in only one significant term (GO:0003676 ‘nucleic acid binding’) shared by *L. sinapis* and *L. reali* (Supplementary table 20), but the same term was marginally non-significant in *L. juvernica* (Fisher test, p = 0.055).

## 4 DISCUSSION

Gene expression is generally under stabilizing selection (Bedford and Hartl, 2009), since altered expression can result in deleterious effects (Wang and Zhang, 2011). At the same time, the transcriptome shows a remarkable redundancy and buffering capacity (Kusnadi et al., 2022), and adaptive evolution of gene expression can sometimes occur rapidly (Hamann et al., 2021). It is therefore not surprising that dosage compensation has evolved independently multiple times across eukaryotes with distinct sex chromosomes (Gu and Walters, 2017). However, the long-held view that dosage compensation mainly restores the expression levels of the proto-sex chromosomes, is compromised by recent findings of mechanisms that rather decrease the expression balance between the sex chromosomes and the autosomes (Mank, 2013). The knowledge about processes that give rise to different modes of dosage compensation is still in its infancy, but organisms with recently recruited neo-sex chromosomes provide good systems to investigate this in more detail. Here, we characterized how dosage compensation and / or dosage balance has established in butterflies that have undergone a dramatic genome restructuring and acquired several neo-Z chromosomes with different levels of degeneration of the corresponding W chromosomes (Höök et al., 2023; Yoshido et al., 2020).

The assessment of sex-biased genes revealed expression balance between the sexes in heads, but a considerable amount of sex-biased expression in abdomens. This is expected since the abdomens contain the gonads, and similar patterns have previously been observed in, for example, chicken (Mank and Ellegren, 2009) and *Drosophila* (Vicoso and Bachtrog, 2015). This indicates that dosage compensation operates differently or is absent in gonads, leading to a general expression bias between the sexes (Gu et al., 2017). Our analysis of dosage compensation / balance on the ancestral and neo-Z chromosomes in *Leptidea* revealed rather complex patterns, but some general trends were noted. For most homologous Z chromosome regions, the expression patterns in the different stages and tissues were largely conserved between the species and we will therefore discuss these from the ancestral *Leptidea* karyotype point of view, which is inferred to be conserved in *L. sinapis* (Höök et al., 2023; Yoshido et al., 2020). For the Z chromosome that contains the ancestral Lepidoptera Z chromosome in *Leptidea*, here referred to as Z1, we observed different modes of dosage compensation for the ancestral (Z1a) and the neo-Z (Z1b) parts, respectively. As previously shown in several moths and butterflies (Catalán et al., 2018; Gu et al., 2017, 2019; Höök et al., 2019; Huylmans et al., 2017; Walters et al., 2015), genes located on the ancestral Z chromosome were generally downregulated in males, i.e. not dosage compensated, but dosage balanced between males and females. In contrast, the genes on the neo-Z fragment that is attached to the ancestral Z chromosome in *Leptidea* (Z1b), were upregulated in females and expressed at similar levels as the autosomal genes across all sample groups. This mimics the situation in the monarch (*Danaus plexippus*), where a rather recently recruited neo-Z part, which has fused to the ancestral Z, is completely dosage compensated as a result of upregulation in females (Gu et al., 2019). However, it contrasts with the situation in the codling moth (*Cydia pomonella*), where a fused neo-Z part is downregulated in males and thus shows similar dosage balance to the ancestral Z chromosome (Gu et al., 2017). We found similar patterns of dual dosage compensation mechanisms for the second Z chromosome (Z2) in *Leptidea*, the only chromosome that has completely conserved synteny between the three *Leptidea* species (Höök et al., 2023). Genes located on the older neo-Z region of this chromosome (Z2a), which became sex-linked more than 13 MYA, were generally dosage compensated via upregulation in adult females. Conversely, genes on the younger Z2 region (Z2b), which was recruited after the split between *L. duponcheli* and the other *Leptidea* species ∼ 13 MYA (Wiemers et al., 2020), were consistently downregulated in males and thus showed dosage balance between the sexes, but overall reduced expression compared to autosomal genes. Hence, both Z1 and Z2 have dichotomous patterns of dosage compensation / balance in *Leptidea*. Our data did not allow for more detailed investigations of underlying mechanisms, but distinct histone modifications in the differently compensated / balanced Z regions, as observed in *D. plexippus* (Gu et al., 2019), could potentially be at play.

It is obviously difficult to dissect the ultimate forces leading to differences in dosage compensation and / or dosage balance between different Z chromosome regions. The situation for Z1 and Z2 in *Leptidea* is analogous to the different modes observed for the ancestral- and neo-Z regions in *D. plexippus* (Gu et al., 2019), but differs from the situation in *C. pomonella* (Gu et al., 2017). This variation warrants further taxonomic sampling to give a more complete picture of the different modes that operate across taxa with different types of neo-sex chromosomes and stands in contrast with the stipulated expectations of co-option of ancestral and neo-sex chromosome (Marín et al., 1996). Here, Lepidoptera might constitute a key system, since independent autosome-Z chromosome fusions have occurred relatively frequently in this group (Wright et al., 2024). Intuitively, the ‘complete dosage compensation’ that occurs for genes on the two older neo-Z regions via upregulation in *Leptidea* females is unlikely just a temporary step in the development towards the same mode of dosage balancing that is operating on the ancestral Z chromosome in many lepidopterans (Catalán et al., 2018; Gu et al., 2017, 2019; Höök et al., 2019; Huylmans et al., 2017; Walters et al., 2015), since the more recently recruited Z2b region actually showed such dosage balance between males and females. The functional categories of the genes located on the autosomal regions that become neo-Z chromosomes could potentially influence the predisposition of different neo-Z regions to develop dosage balance or dosage compensation. We found some GO term enrichments in the regions with different modes in *Leptidea*, but each term was represented by very few genes and inferring an association between specific gene functions and modes of regulation therefore becomes speculative. A general explanation could be that neo-Z chromosome regions that become dosage compensated (i.e. female upregulation of the Z-linked genes) contain a higher proportion of ‘dose sensitive’ genes, which impose stronger selection to maintain the ancestral expression levels, while neo-Z regions that evolve towards a dosage balance are less sensitive to dose differences between Z-linked and autosomal genes. An indication in this direction is that we found that sulfate transport genes, which have been shown to be haploinsufficient in mice (Zhang et al., 2019), were only present in regions that were dosage compensated in *Leptidea*. Characterizing dosage sensitivity for all genes located on neo-sex chromosomes will therefore be an important step to understand the forces leading to different types of compensating or balancing mechanisms.

Previous analysis based on hybridization with a handful of probes have indicated that the most recently derived W chromosome (W3) in *Leptidea* contains some functional genes (Yoshido et al., 2020). By using female-specific polymorphisms we could verify that W3 harbors at least 56 single copy gametologs that are present in all the three *Leptidea* species analyzed here. Note that we did not find strong evidence for functional gametologs on the other sex chromosomes. Of the 56 identified W3-linked gametologs, 39-54% were expressed in females in the different species and the overall expression patterns observed for the gametologs on the homologous neo-Z chromosome (Z3) suggested that genes on this chromosome were at least partly dosage compensated via upregulation in females. However, a much more complex pattern was revealed when analyzing the relative expression of the Z3- and W3-linked gametologs across all species. Specifically, in head tissue, the Z3 / W3 gametologs had significantly higher allelic expression imbalance in females than the Z3-linked gametologs in males. This indicates that W3-linked alleles are specifically regulated or that degeneration is affecting the expression of a subset of W3-linked gametologs and shows that dosage compensation has not yet evolved for these genes. Conversely, we found no expression imbalance between the Z / W-pair alleles in adult female abdomen, which suggests that tissue-specific regulation occurs. This likely reflects the obvious differences in tissue composition between head and abdomen and / or that trans regulation is more common in abdominal tissue. We also found that the genes with bi-allelic expression (Z3 and W3 gametolog expressed) in females had significantly higher expression levels than autosomal genes in both males and females. It is tempting to speculate that this set of genes was highly expressed already on the proto-Z3 chromosome and that these particular genes are sensitive to changes in both relative dose and absolute expression levels, i.e. that expression has been maintained at comparatively high level in both males in females. Interestingly, genes with mono-allelic expression in females (Z / 0) had reduced expression compared to the autosomes in both males and females, which shows that males downregulate these genes specifically. These results show that inter-sexual dosage balance has evolved rapidly on Z3 and occurs on a gene-by-gene basis (or in smaller genomic blocks) on the relatively recently recruited neo-Z3 chromosome.

In Lepidoptera, there is female achiasmy (Turner and Sheppard, 1975), which means that neo-W chromosomes immediately stop recombining. Consequently, a comparatively rapid degeneration, with loss of functional genes and enrichment of repetitive elements, is predicted (Shipilina et al., 2022; Traut et al., 2013). It might therefore be surprising that we identified *>* 50 actively expressed, and apparently functional, W3-linked genes in the three focal *Leptidea* species. However, the W3-linked genes had high expression levels in general, which indicates that they are under considerable selection pressure to maintain function (Charlesworth, 2021). We also know that deleterious effects of selection on linked sites should decrease with the loss of functional genes on the W chromosome (Charlesworth, 2021), and this may have contributed to maintenance of functional genes on the *Leptidea* W3 chromosome. The considerable overlap of retained and expressed W3-linked orthologs across species indicates that gene loss predominantly occurred prior to the split of the three species analyzed here. This state is analogous to the evolution of the neo-Y chromosome in *Drosophila miranda*, where male achiasmy has led to rapid, but not complete, degeneration of genes since the recruitment of this chromosome ∼ 1.5 MYA (Bachtrog et al., 2008). One observed difference between these two systems relates to retainment of specific functional categories; no specific functional classes are overrepresented among the still expressed genes on the *D. miranda* neo-Y (Bachtrog et al., 2008) while genes associated with nucleic acid binding were enriched among Z3-linked genes with expressed W3 gametologs in *Leptidea*. This suggests that these specific genes are under functional constraint on the W3 chromosome in *Leptidea*, but it should also be noted that a relatively high number of genes in this functional category appear to have lost their W-linked gametolog.

In theory, the distinctive expression patterns observed for the different Z-linked regions could reflect retained variation in ancestral autosomal expression levels rather than variation associated with the evolution of dosage balance / compensation. However, a simple visual inspection shows that individual autosomes vary only marginally in expression levels compared to the genome wide average (Supplementary figure 10). In addition, there is no consistent trend across stages and tissues as observed for e.g. the ancestral Z chromosome and Z2b. Explicit modelling of expression evolution of orthologs that are sex-linked in *Leptidea*, but located on autosomes in a set of outgroup species could indicate if the inferred ancestral expression levels are expected to deviate from the neutral expectation for this particular group of genes, but such an analysis would require data that we do not have access to.

In contrast to comparative approaches between deeply divergent lineages, analyses of gene regulation in species with multiple sex chromosomes can control for lineage-specific effects on the evolution of dosage compensation. If species-specific traits or characteristics are important for establishment of certain dosage compensation / balance mechanisms, analogous mechanisms would be expected to evolve repeatedly in organisms with similar characteristics. Here, we show that different modes of regulation have evolved in a single lineage, which suggests that additional factors are important for how dosage regulation is established and maintained. However, it is important to recognize that several properties may be different in butterflies compared to other systems. Effective population sizes should for example be considerably larger in butterflies (Mackintosh et al., 2019; Talla et al., 2019; Boman et al., 2025) compared to other female heterogametic systems like birds and snakes, where dosage compensation is mostly incomplete (Ellegren et al., 2007; Vicoso et al., 2013). Such differences in effective population size will manifest in differences in the efficacy of selection and the propensity for specific mechanisms to evolve. Our observations in *Leptidea* highlight the complexity of gene dosage evolution on sex chromosomes and show that neo-sex chromosomes are not bound to be co-opted with the mechanism that are already present on the ancestral sex chromosome.

## 5 DATA AVAILABILITY

All raw sequencing reads have been deposited at the European Nucleotide Archive under accession *PRJEB66419*. All scripts used in the analysis are available at Github (https://github.com/EBC-butterfly-genomics-team/Leptidea_Z_chr_expression).

## 6 ACKNOWLEDGEMENTS

This work was funded by the Swedish Research Council (VR research grant #019-04791 to N.B.), NBIS/SciLifeLab long-term bioinformatics support (WABI) and The Swedish Collegium for Advanced Science (Natural Sciences Programme, Knut and Alice Wallenberg Foundation, Postdoc funding for D.S.). The authors acknowledge support from the National Genomics Infrastructure in Stockholm funded by Science for Life Laboratory, the Knut and Alice Wallenberg Foundation and the Swedish Research Council, and SNIC/Uppsala Multidisciplinary Center for Advanced Computational Science for assistance with massively parallel sequencing and access to the UPPMAX computational infrastructure. The computations were enabled by resources provided by the National Academic Infrastructure for Supercomputing in Sweden (NAISS) and the Swedish National Infrastructure for Computing (SNIC) in Uppsala, partially funded by the Swedish Research Council through grant agreements no. 2022-06725 and no. 2018-05973. R.V. was supported by Grant PID2022-139689NB-I00 (MICIU/AEI/10.13039/501100011033 and ERDF, EU), and by grant 2021-SGR-00420 (Departament de Recerca i Universitats, Generalitat de Catalunya). Travel grants awarded to K.N. and L.H. from Bjurzons resestipendium and to K.N. from Thelins G resestipendium contributed to fieldwork for sample collection. We are grateful for the substantial support and encouragement for the project we received from Professor Christer Wiklund.

## Notes

### Competing Interest Statement

The authors have declared no competing interest.

