## Supplementary Information for "Contrasting Modes of Dosage Compensation on Ancestral and Neo-Z Chromosomes in Butterflies"

#### Supplementary figures

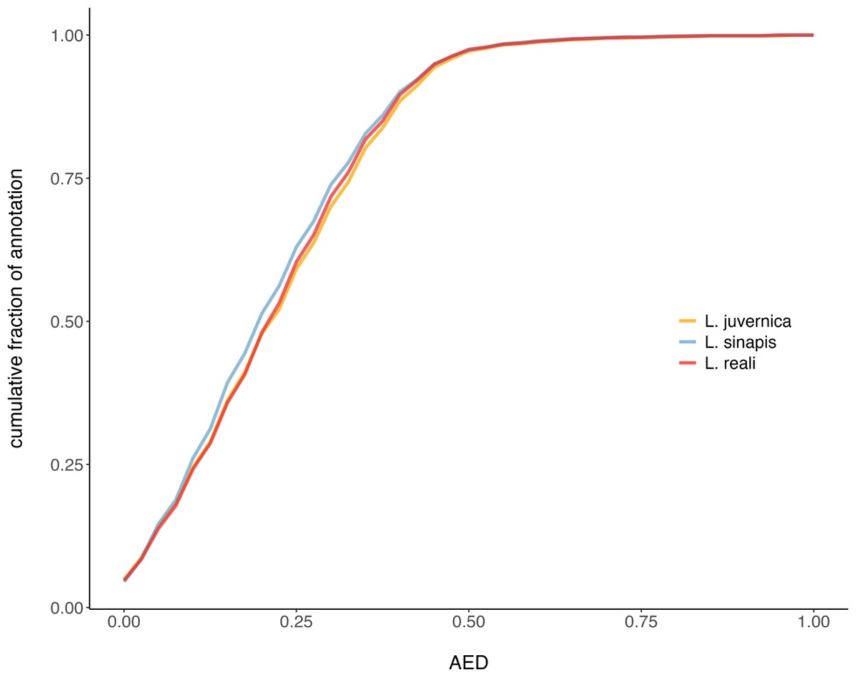

***Supplementary figure 1.*** *Cumulative AED score for gene annotations. An AED score of 0 indicates full congruence and AED = 1 indicates lack of congruence between the evidence and the final annotation.*

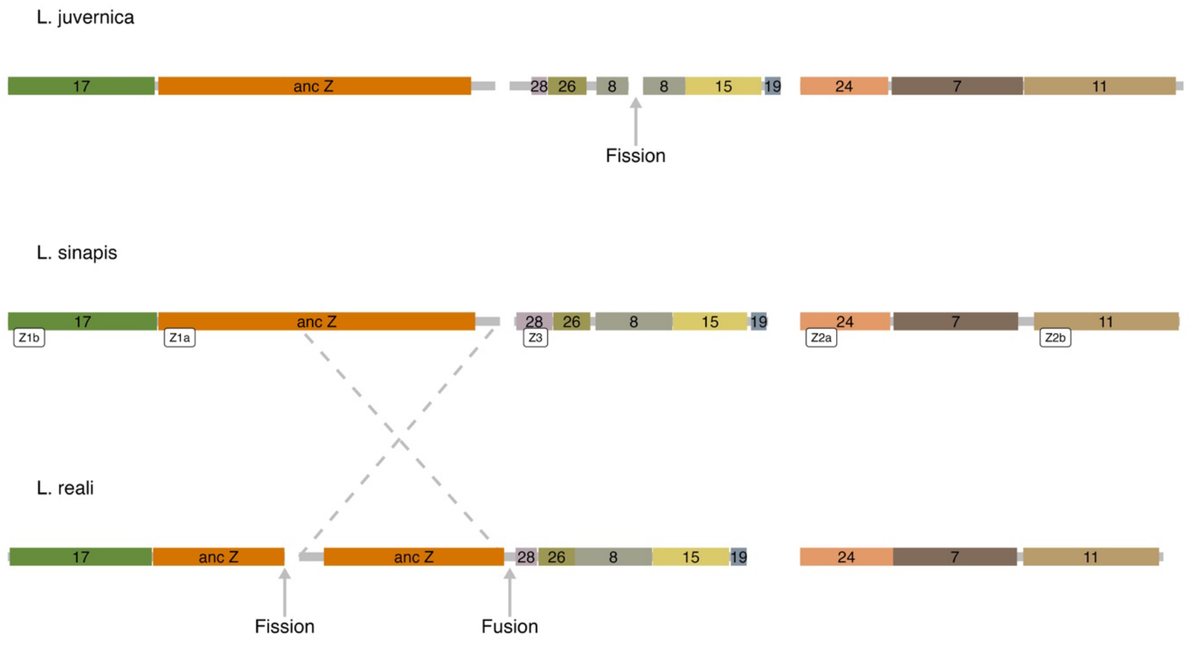

***Supplementary figure 2.*** *Previously inferred syntenic relationships and rearrangements of the Z chromosomes in the three Leptidea species. Since the fission and fusion event in L. reali occurred on the same branch it is possible that they constitute a translocation (one molecular event) rather than a fission followed by a fusion. The numbers on the chromosomes (colored blocks) indicate homologous regions of autosomal fragments in B. mori. The chromosomes are ordered and oriented to enhance visualization. The dashed lines indicate a region that is plotted with inverted orientation in L. reali relative to the other species. The white labels under each Z chromosome in L. sinapis show regions that were analyzed separately in this study.*

A

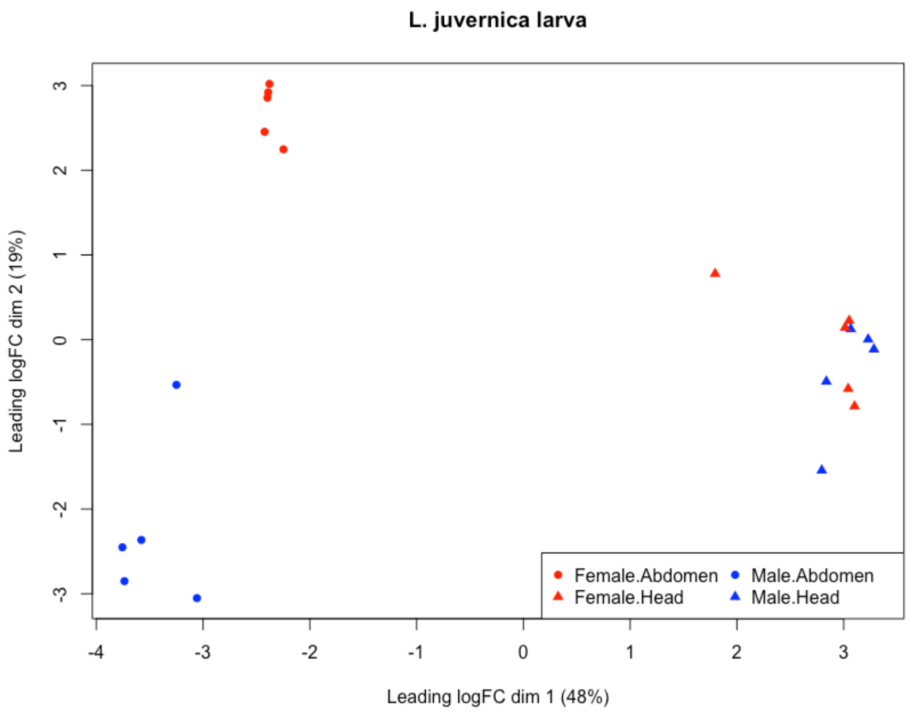
L. juvernica larva
Leading logFC dim 2 (19%)
3
2
1
0
-1
-2
-3
-4
-3
-2
-1
0
1
2
3
Leading logFC dim 1 (48%)
Female.Abdomen
Female.Head
Male.Abdomen
Male.Head

B

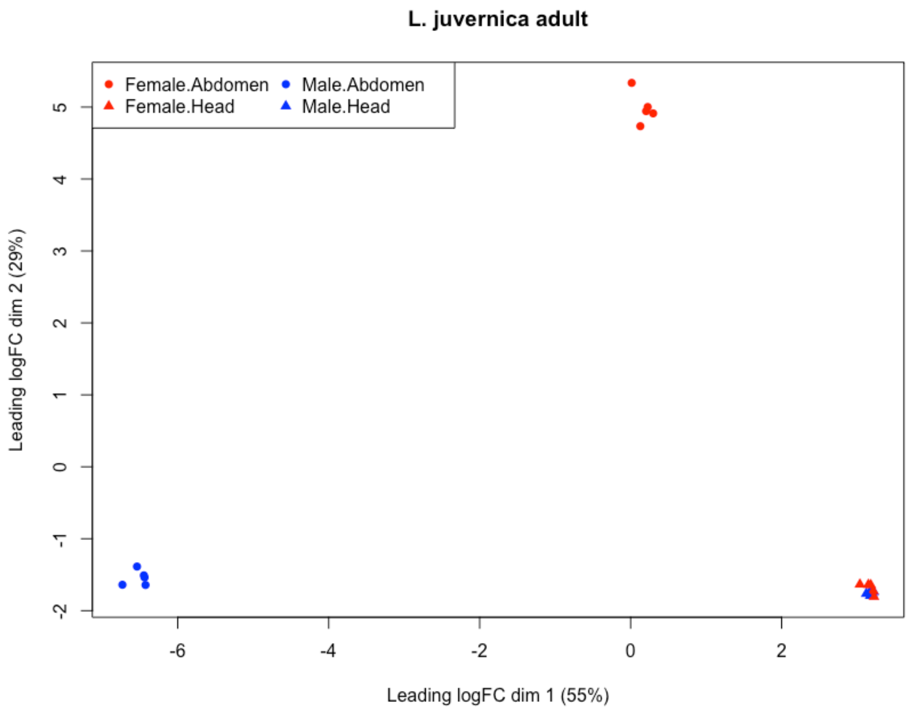
L. juvernica adult
Leading logFC dim 2 (29%)
5
4
3
2
1
0
-1
-2
-6
-4
-2
0
2
Leading logFC dim 1 (55%)
Female.Abdomen
Female.Head
Male.Abdomen
Male.Head

C

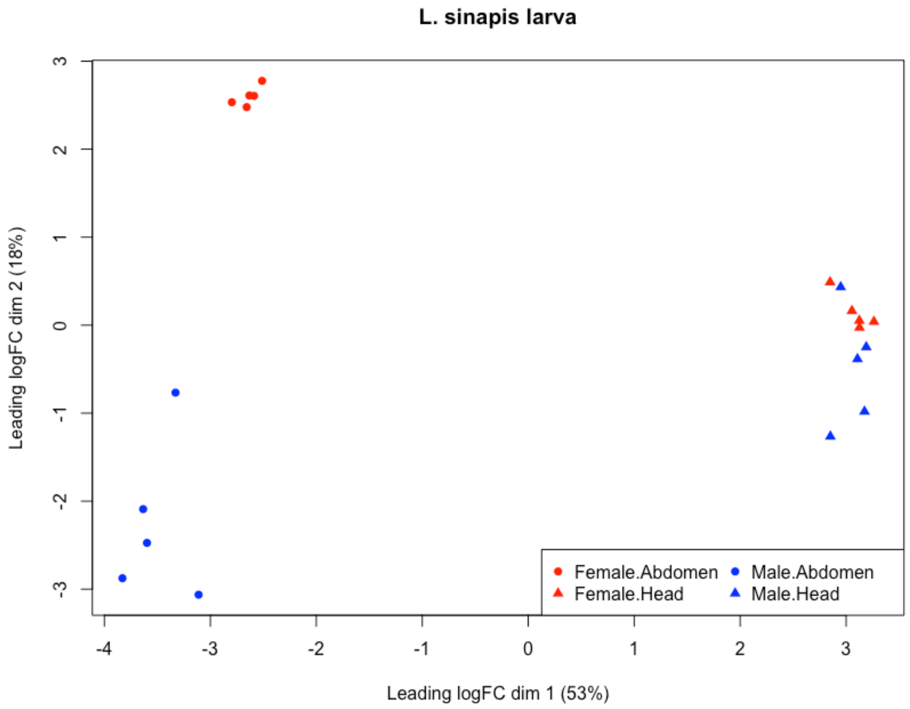
L. sinapis larva
Leading logFC dim 2 (18%)
Leading logFC dim 1 (53%)
Female.Abdomen
Female.Head
Male.Abdomen
Male.Head

D

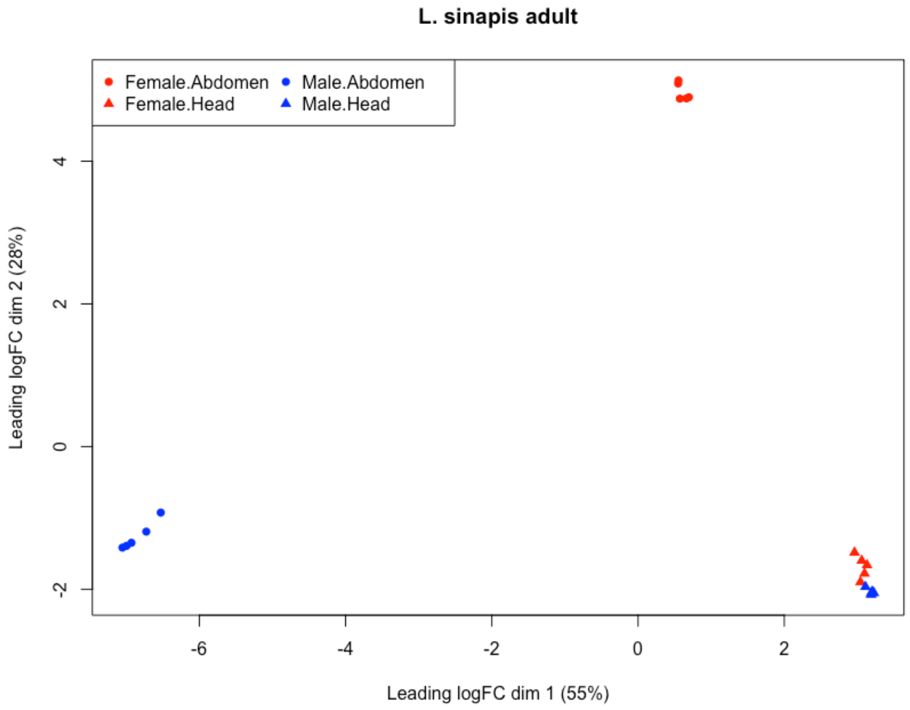
L. sinapis adult
Leading logFC dim 2 (28%)
Leading logFC dim 1 (55%)
Female.Abdomen
Female.Head
Male.Abdomen
Male.Head

**E**

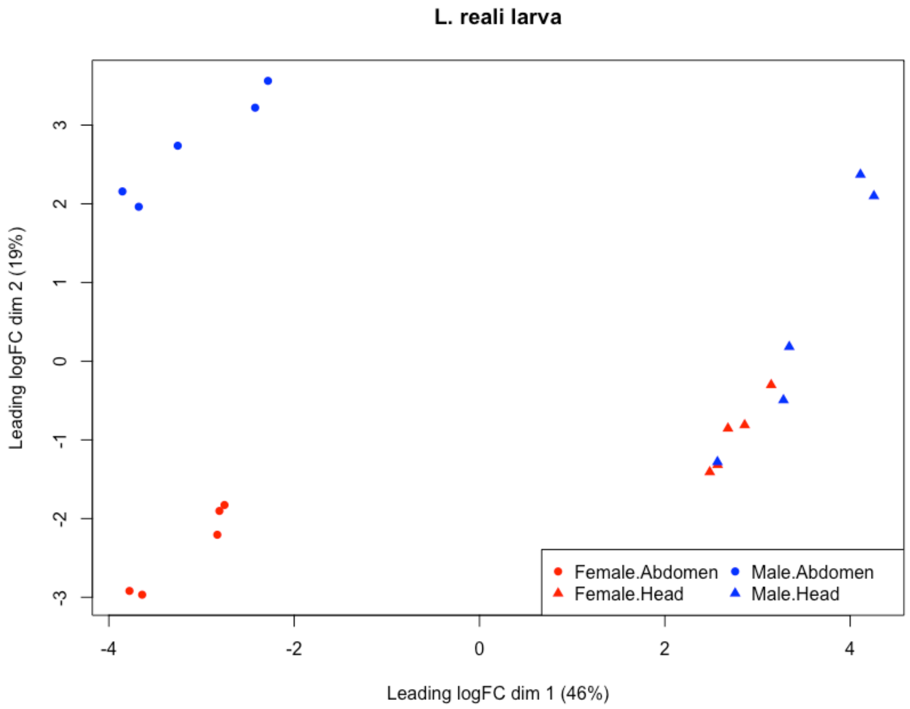

**F**

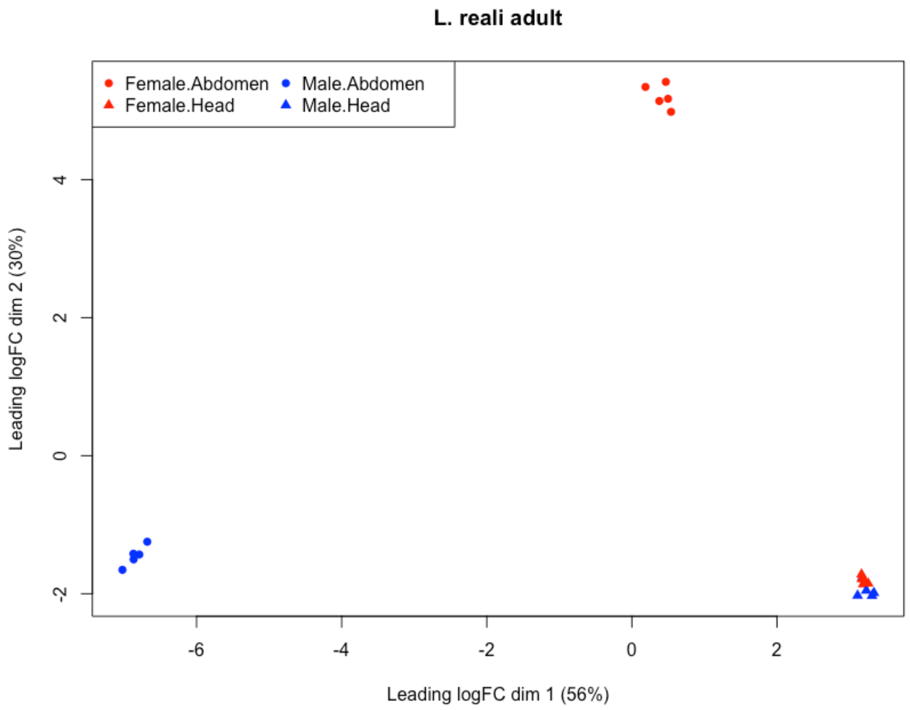

***Supplementary figure 3****. Multidimensional scaling plots showing clustering of samples by sex and tissue based on gene expression in each stage for L. juvernica (A, B), L. sinapis (C, D) and L. reali (E, F).*

A: L. juvernica

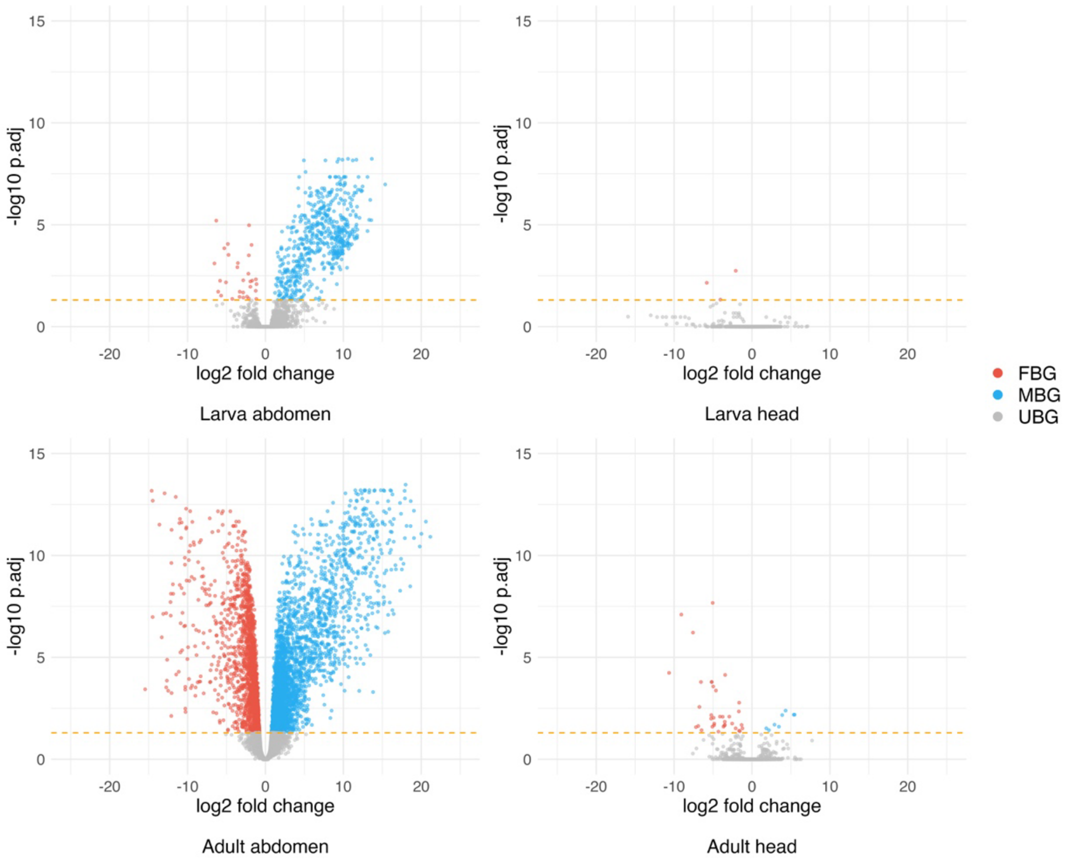

B: L. sinapis

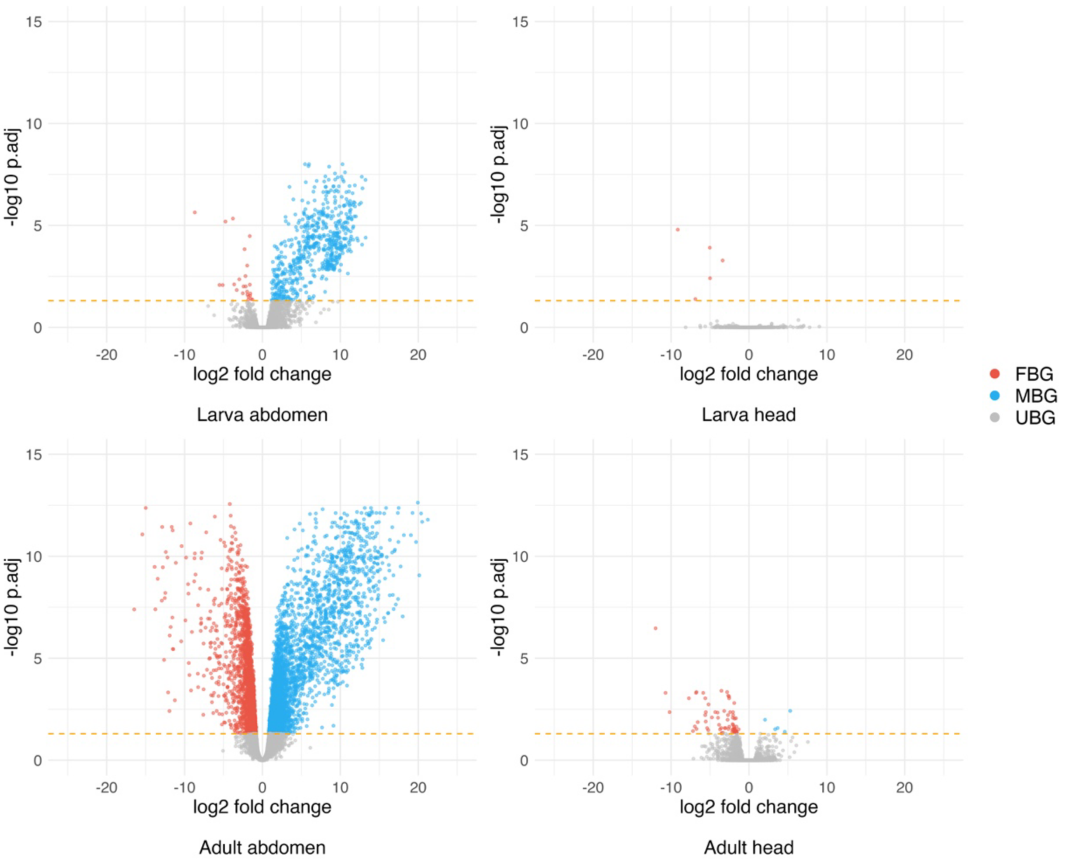
-log10 p.adj
log2 fold change
Larva abdomen
Larva head
Adult abdomen
Adult head
FBG
MBG
UBG

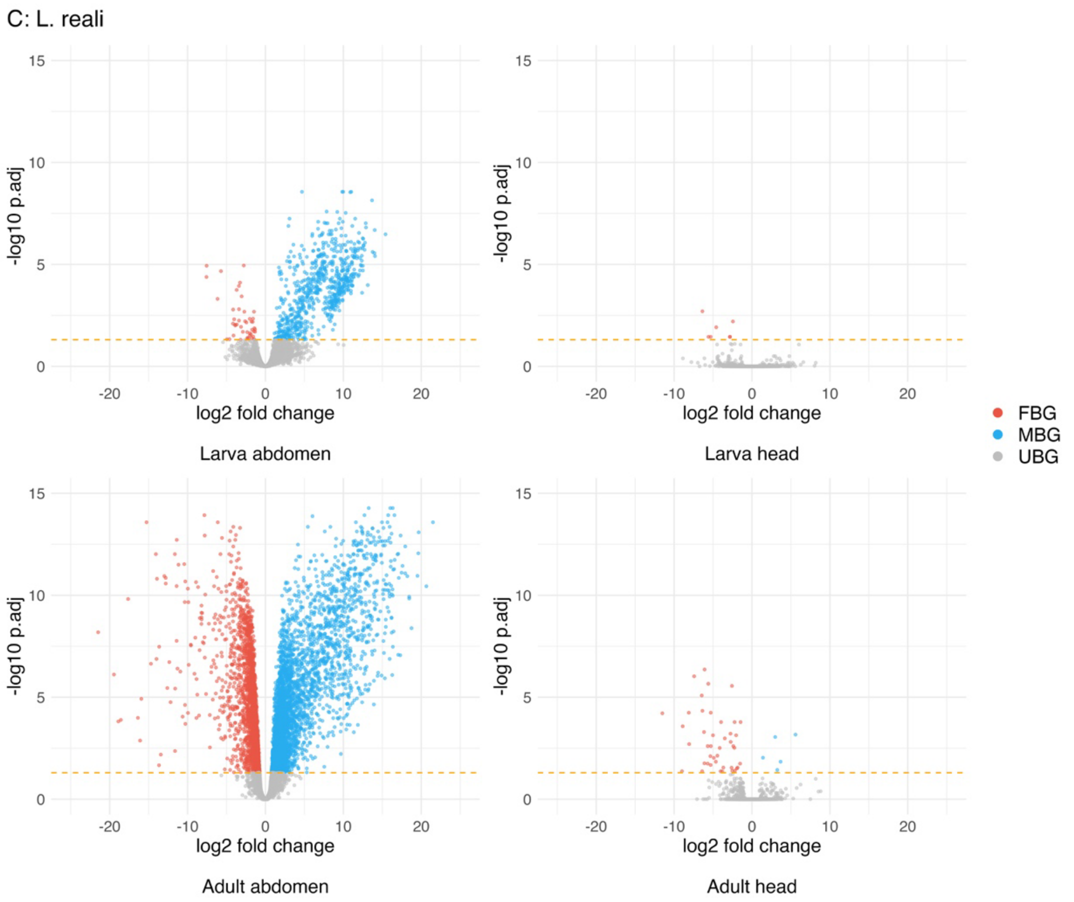

***Supplementary figure 4.*** *Volcano plots showing results from the differential expression analysis between sexes in A, L. juvernica B, L. sinapis and C, L. reali. The dashed lines show the 0.05 FDR cutoff used to define significantly differentially expressed genes, plotted on a -log10 scale. FBG = female-biased genes, MBG = male-biased genes and UBG = unbiased genes.*

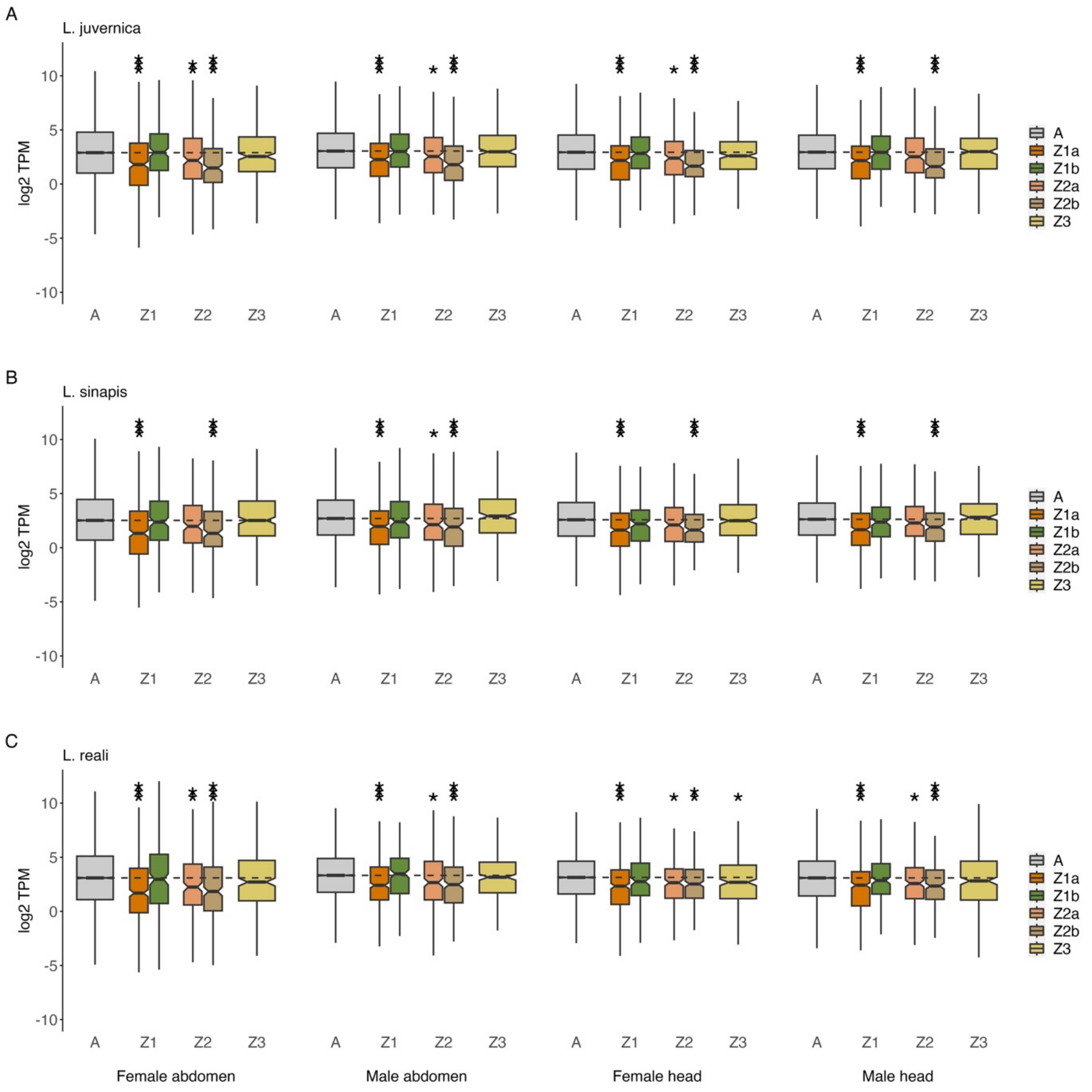

***Supplementary figure 5.*** *Boxplots showing gene expression levels in the larval stage in A, L. juvernica; B: L. reali, and C: L. sinapis. Asterisks show Z chromosome regions that have significantly reduced expression compared to the autosomes (* $P \leq 0.05$, ** $P \leq 0.01$, *** $P \leq 0.001$, Benjamini-Yekutieli corrected).*

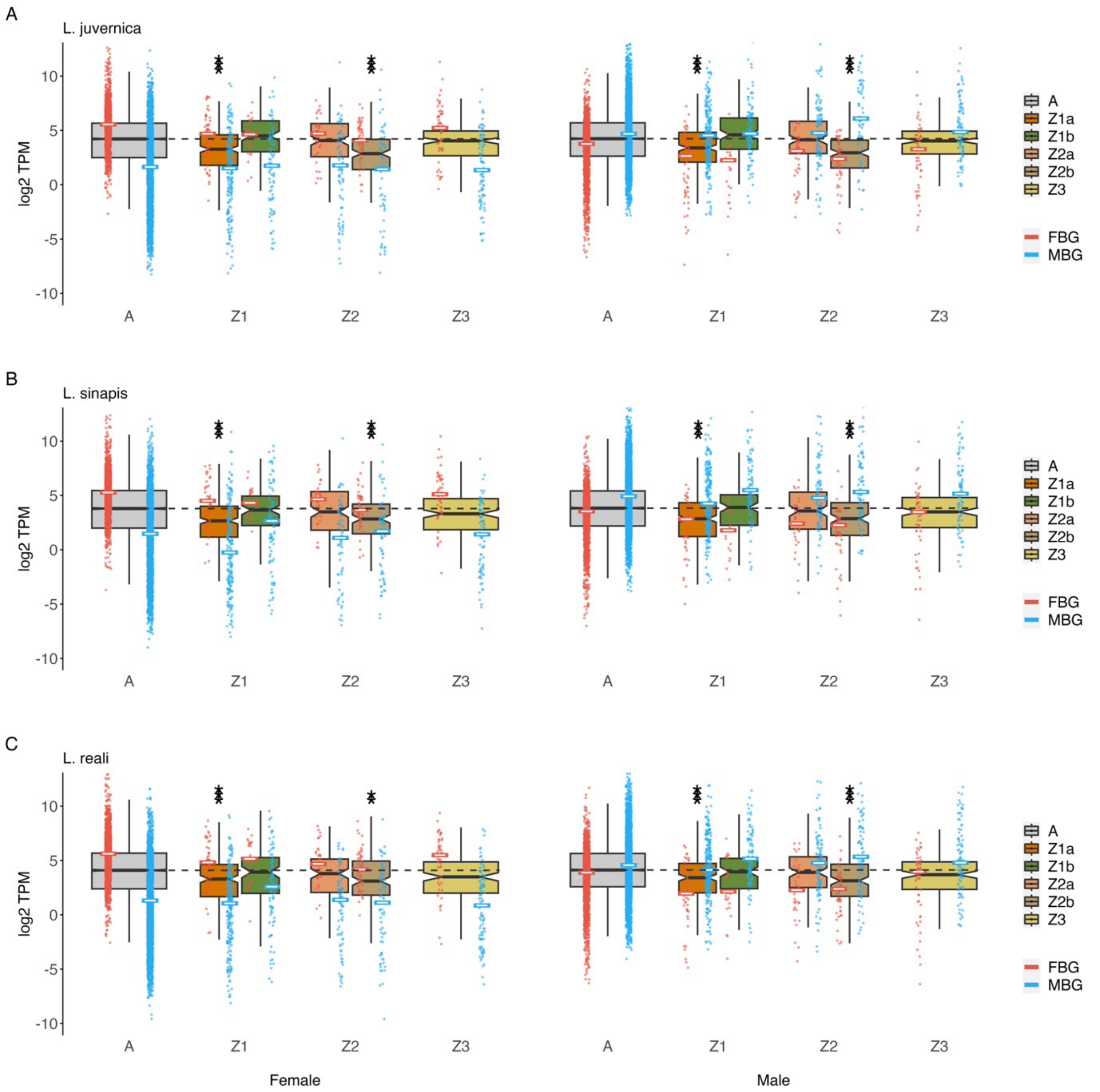

***Supplementary figure 6.*** *Boxplots showing expression levels of unbiased genes in adult abdomen in A, L. juvernica; B: L. reali, and C: L. sinapis. Dots show expression levels of sex-biased genes in each sex and medians are indicated with horizontal lines. Black dotted lines show autosomal median expression levels for each sample group. Asterisks show Z chromosome regions that have significantly reduced expression levels compared to the autosomes (* P $\leq$ 0.05, ** P $\leq$ 0.01, *** P $\leq$ 0.001, Benjamini-Yekutieli corrected).*

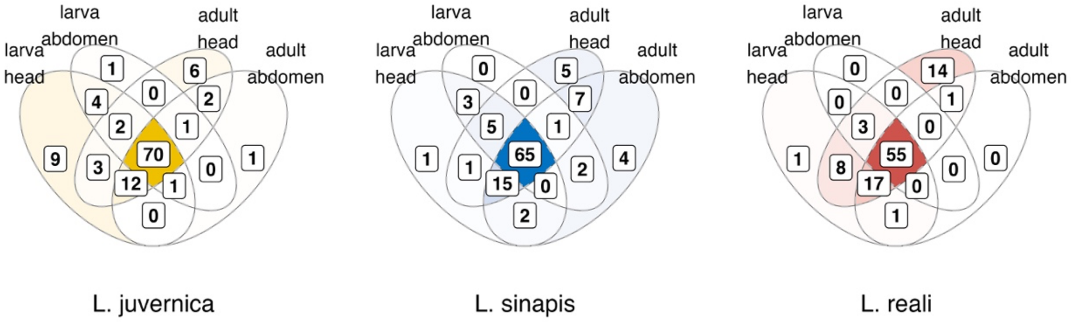

***Supplementary figure 7***. *Overlap of genes with female unique SNPs on Z3 across different stages and tissues in each Leptidea species. Color intensity reflects the number of genes. The count shows the number of genes after filtering for low read counts for both alleles (> 10).*

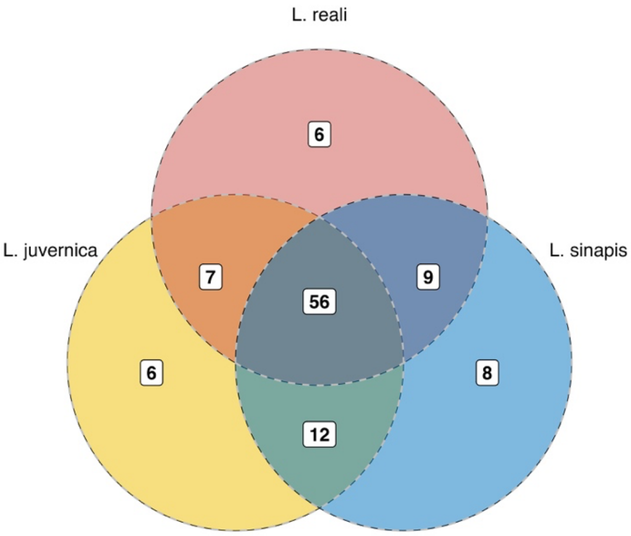

***Supplementary figure 8***. *Overlap of orthologs on Z3 with expression of female unique SNPs. The counts include the union of genes expressed in any stage and tissue.*

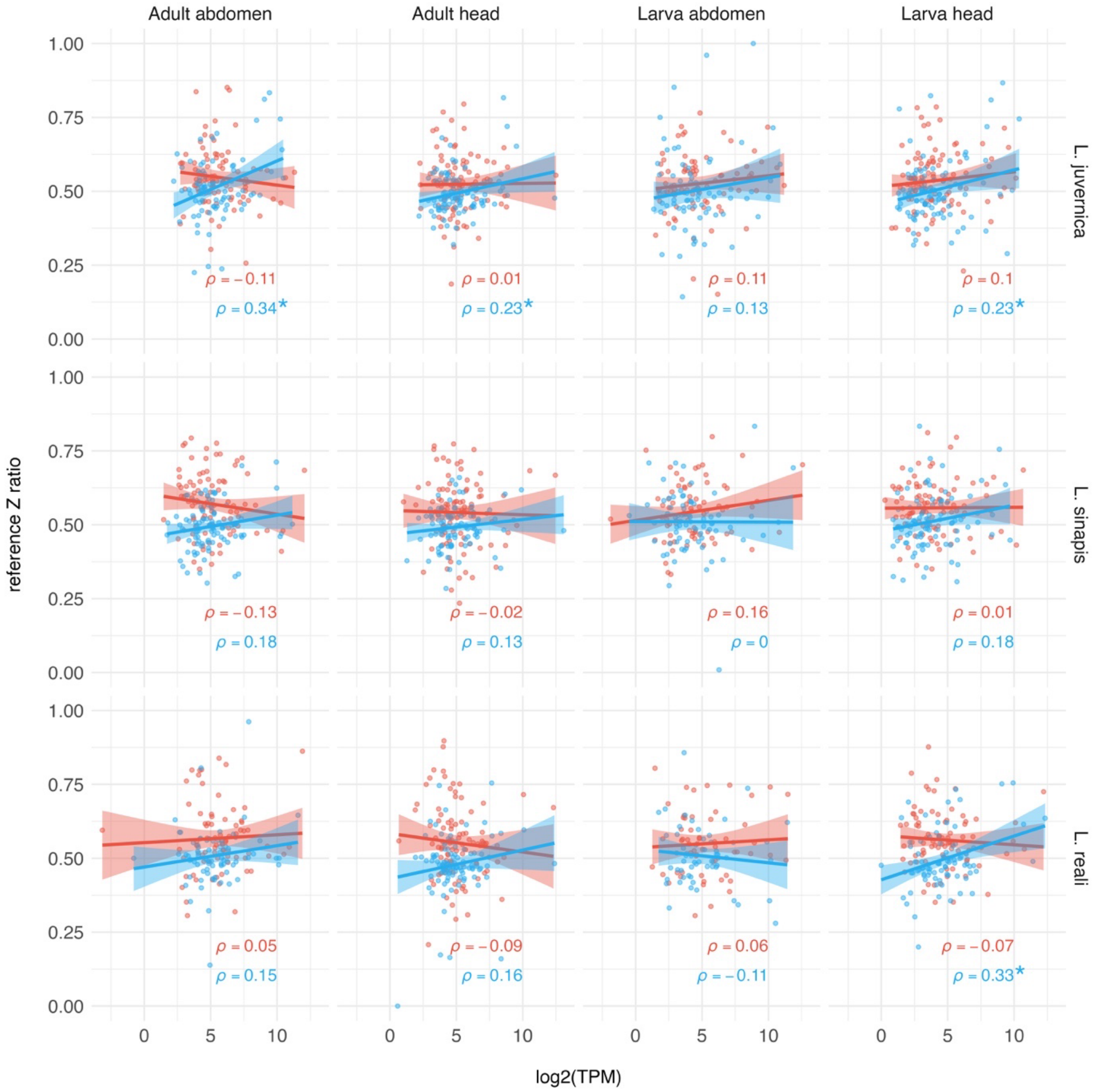

***Supplementary figure 9**. Pearson correlations between the relative expression of the reference Z allele and expression levels (log2 TPM). Significant correlations are indicated with asterisks. Test statistics and p-values are available in **Supplementary table 15**.*

**A**

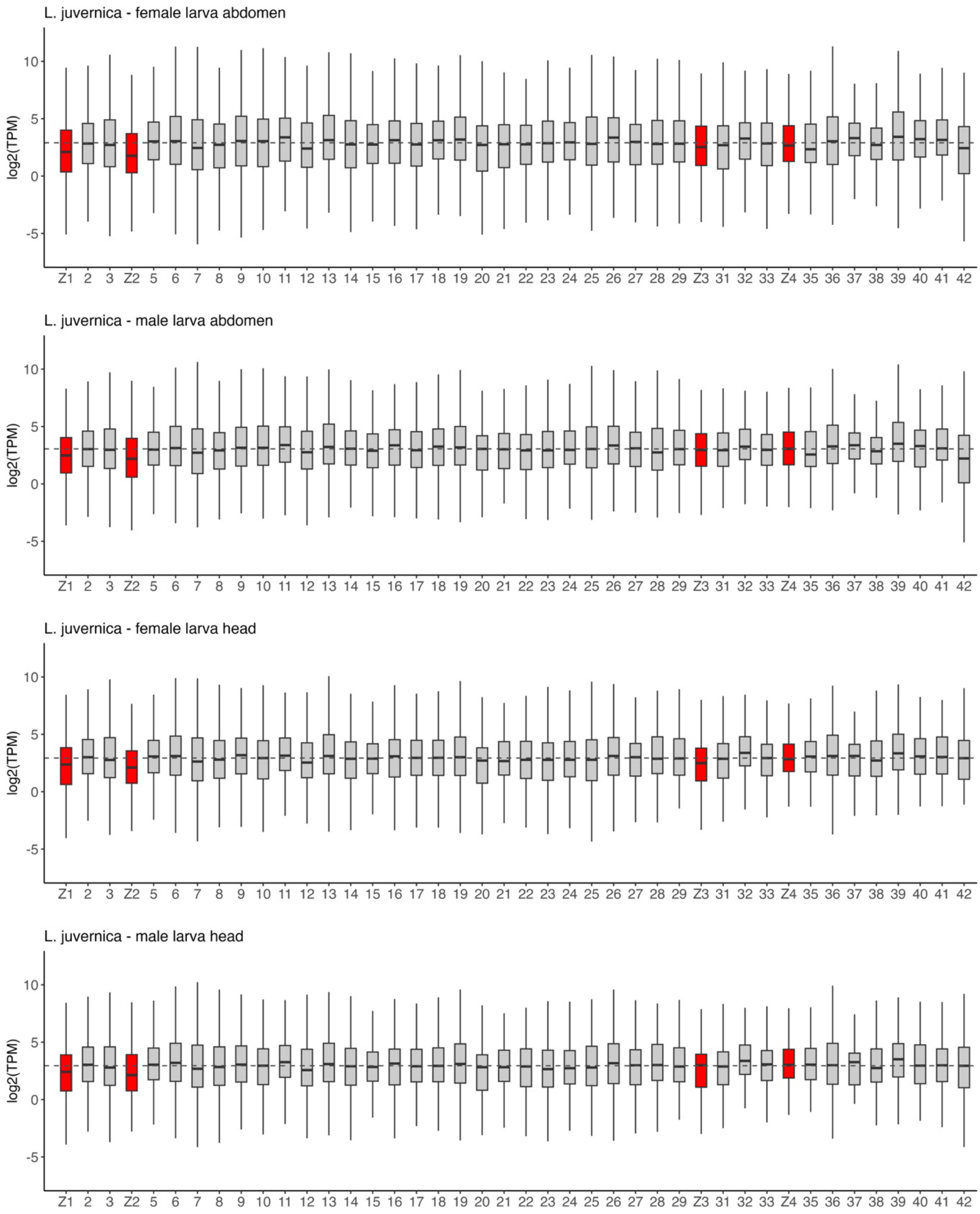
L. juvernica - female larva abdomen
L. juvernica - male larva abdomen
L. juvernica - female larva head
L. juvernica - male larva head
log2(TPM)
10
5
0
-5
Z1 2 3 Z2 5 6 7 8 9 10 11 12 13 14 15 16 17 18 19 20 21 22 23 24 25 26 27 28 29 Z3 31 32 33 Z4 35 36 37 38 39 40 41 42

**B**

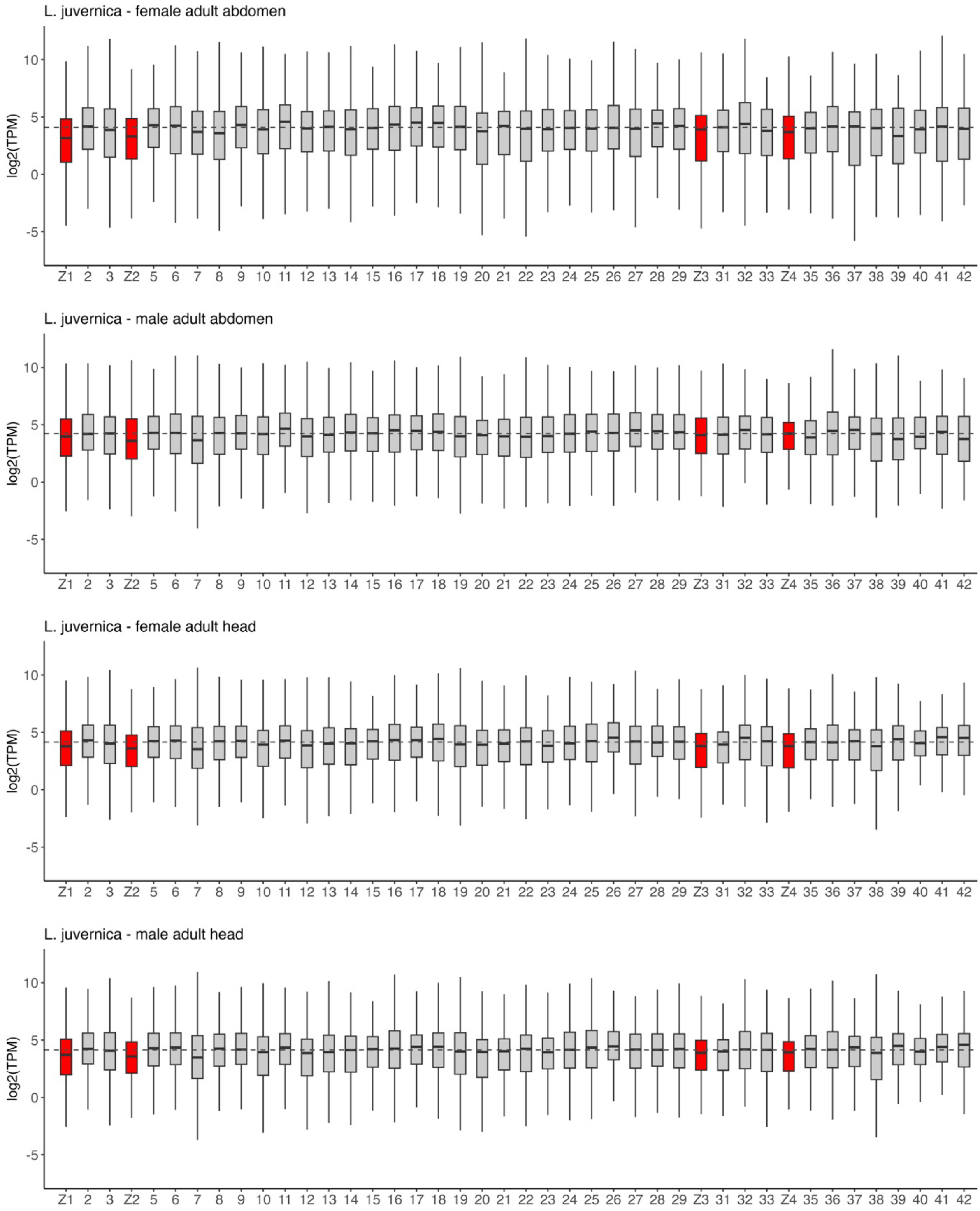
L. juvernica - female adult abdomen
L. juvernica - male adult abdomen
L. juvernica - female adult head
L. juvernica - male adult head
log2(TPM)
10
5
0
-5
Z1 2 3 Z2 5 6 7 8 9 10 11 12 13 14 15 16 17 18 19 20 21 22 23 24 25 26 27 28 29 Z3 31 32 33 Z4 35 36 37 38 39 40 41 42

**C**

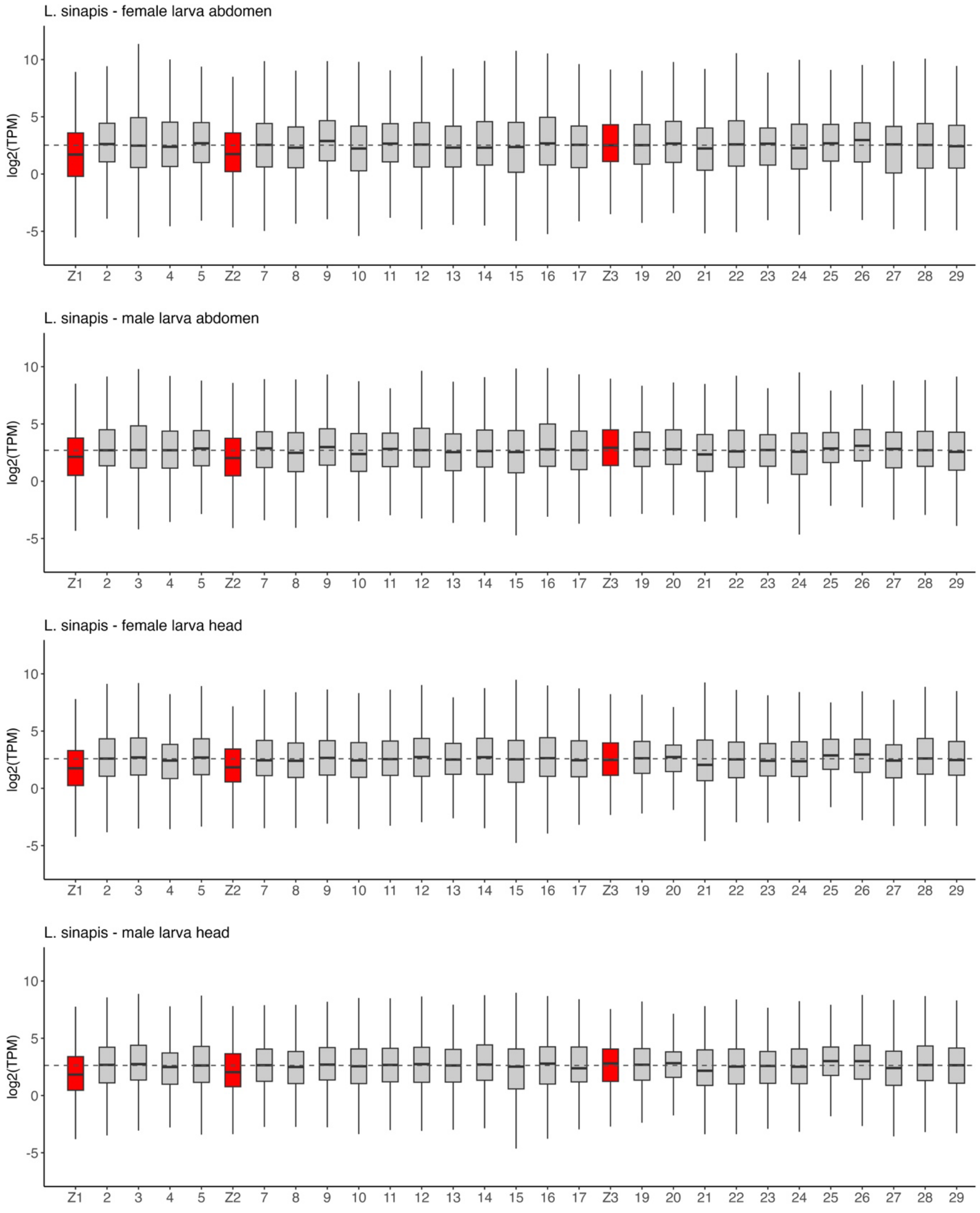
L. sinapis - female larva abdomen
L. sinapis - male larva abdomen
L. sinapis - female larva head
L. sinapis - male larva head
log2(TPM)
10
5
0
-5
Z1 2 3 4 5 Z2 7 8 9 10 11 12 13 14 15 16 17 Z3 19 20 21 22 23 24 25 26 27 28 29

D

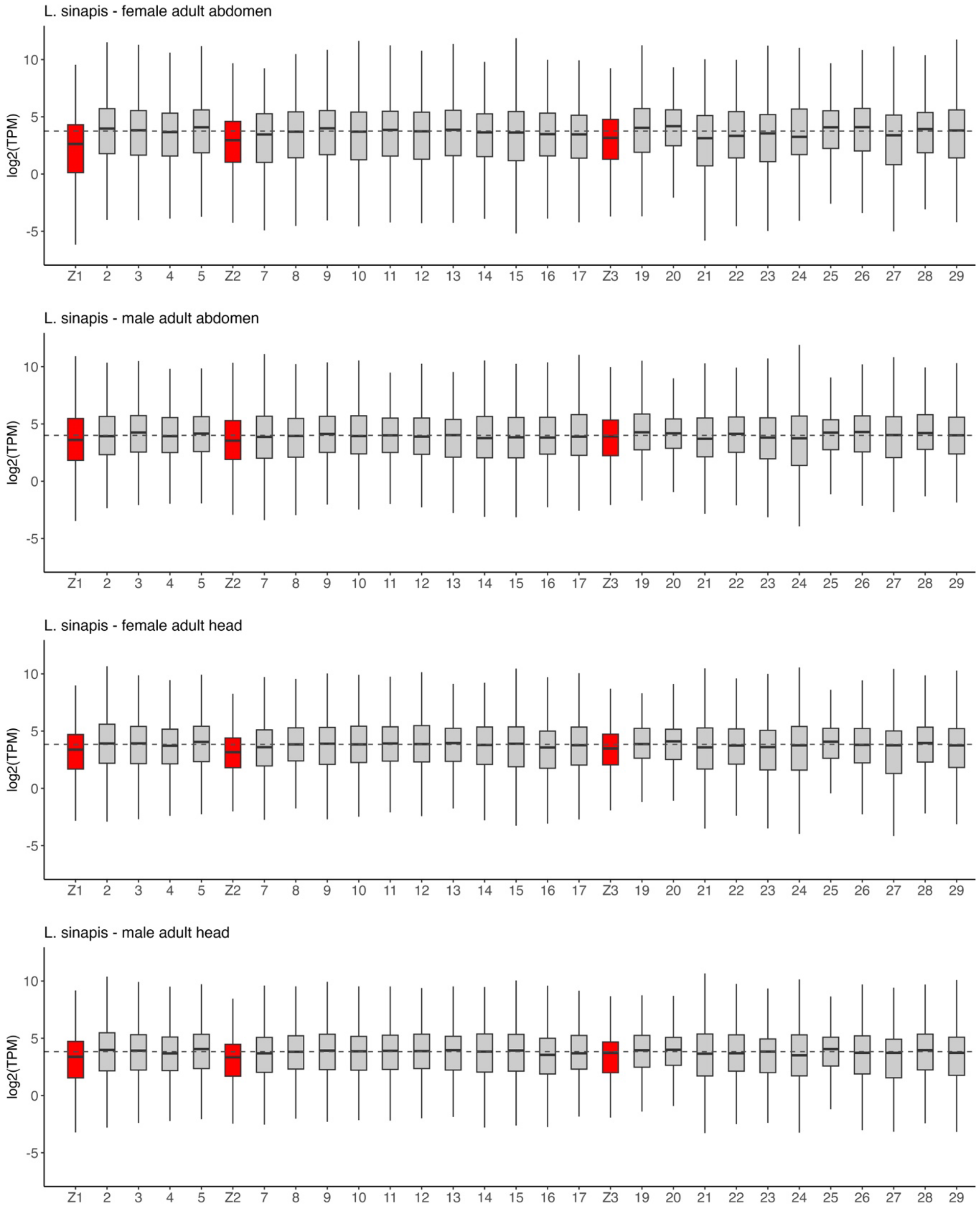

L. sinapis - female adult abdomen
L. sinapis - male adult abdomen
L. sinapis - female adult head
L. sinapis - male adult head
log2(TPM)
10
5
0
-5
Z1 2 3 4 5 Z2 7 8 9 10 11 12 13 14 15 16 17 Z3 19 20 21 22 23 24 25 26 27 28 29

E

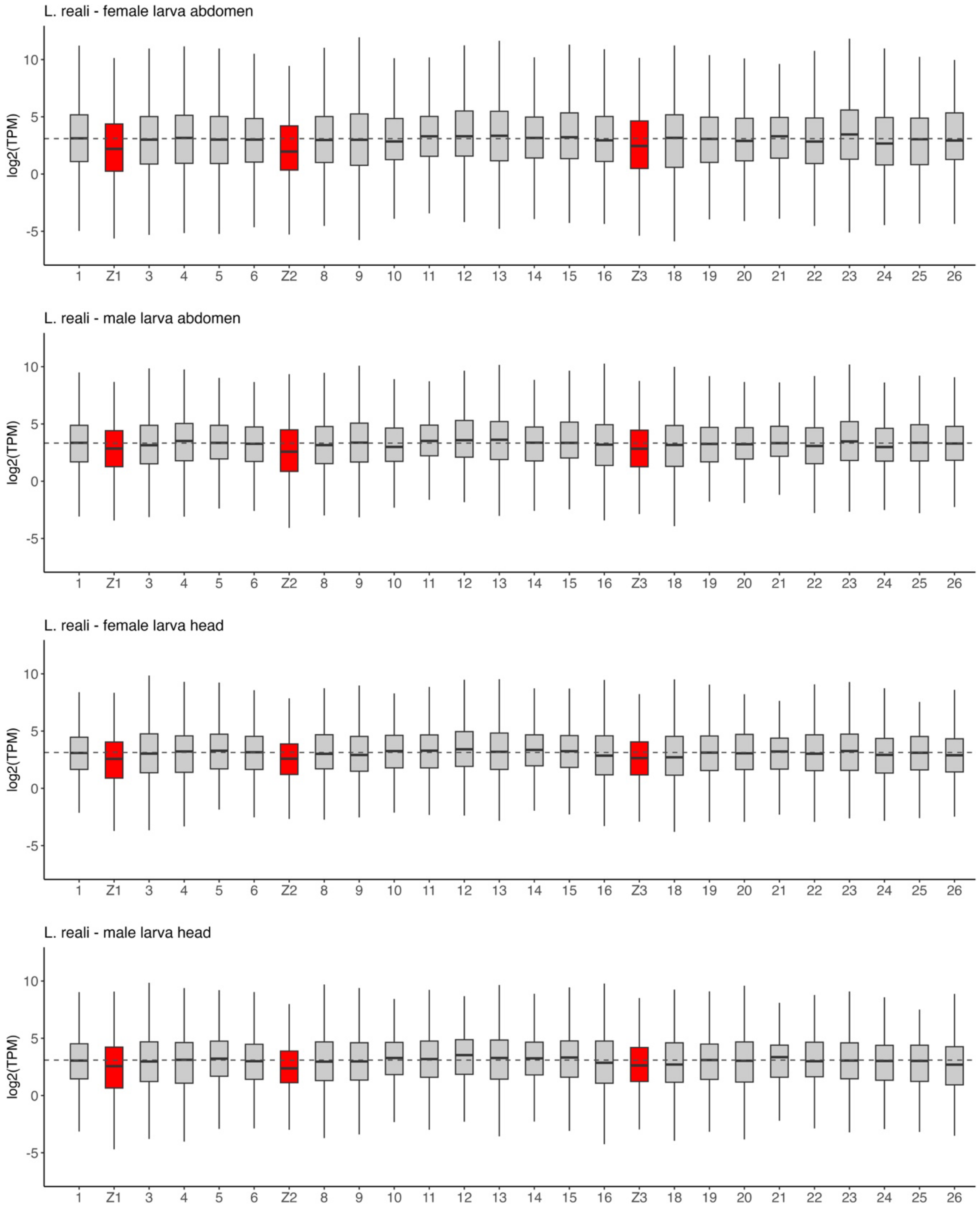

L. reali - female larva abdomen
L. reali - male larva abdomen
L. reali - female larva head
L. reali - male larva head
log2(TPM)
10
5
0
-5
1
Z1
3
4
5
6
Z2
8
9
10
11
12
13
14
15
16
Z3
18
19
20
21
22
23
24
25
26

F

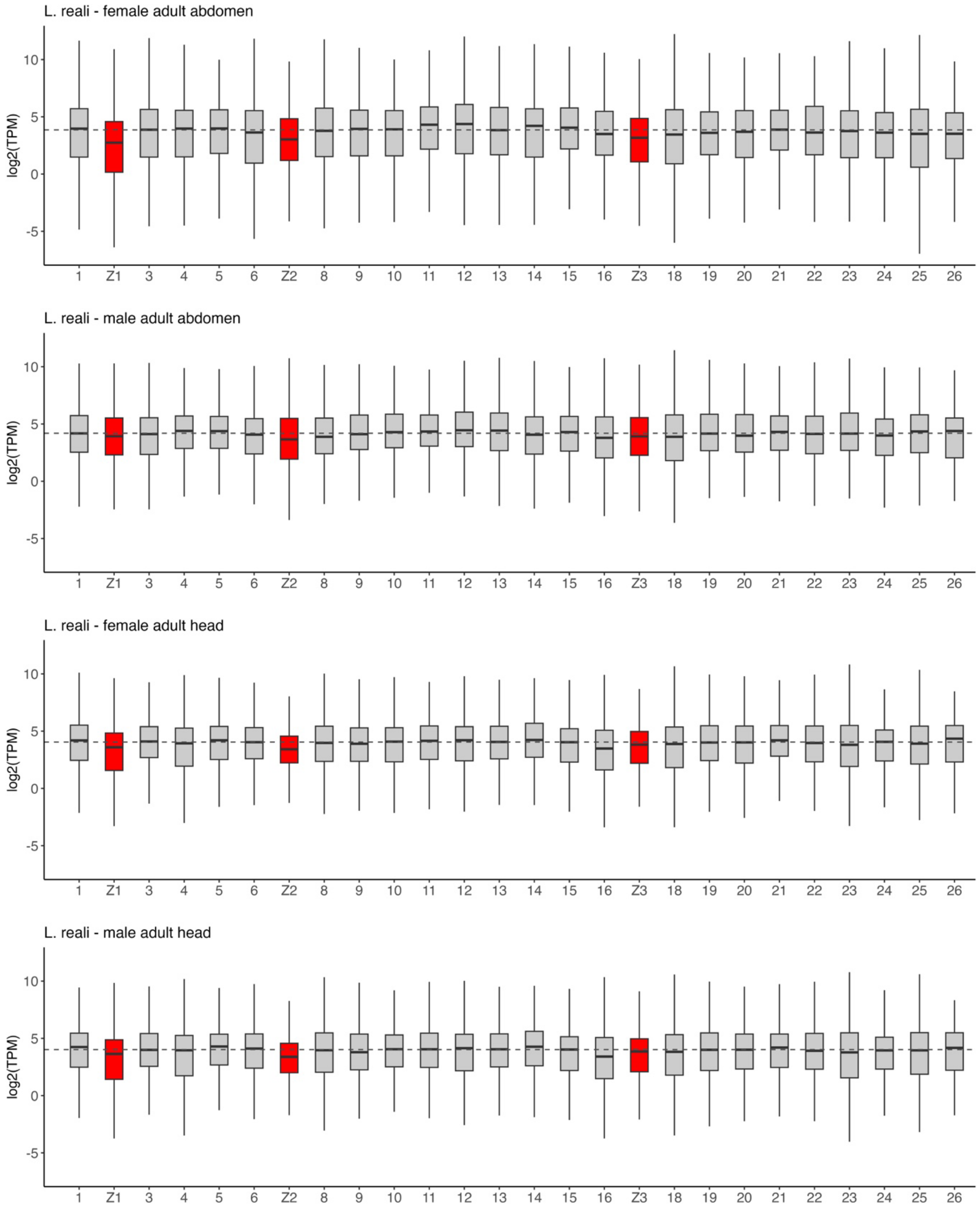

**Supplementary figure 10.** Boxplots showing expression levels of genes on individual chromosomes in A-B, *L. juvernica*; C-D: *L. sinapis* and E-F: *L. reali*. Dashed lines show median autosomal expression for each respective sample group.

### Supplementary tables

**Supplementary table 1.** Result of BUSCO analysis for transcriptomes using the 'lepidoptera_odb10' lineage set containing 5,286 BUSCO groups.

| Species | Sample group | Comp-lete | Complete and single-copy | Complete and duplicated | Frag-mented | Miss-ing |
|---|---|---|---|---|---|---|
| *L. juvernica* | Adult female | 5,028 | 1,492 | 3,536 | 58 | 200 |
| | Adult male | 5,143 | 1,865 | 3,278 | 50 | 93 |
| | Larva female | 4,827 | 2,227 | 2,600 | 82 | 377 |
| | Larva male | 5,029 | 2,292 | 2,737 | 62 | 195 |
| *L. sinapis* | Adult female | 5,014 | 1,911 | 3,103 | 73 | 199 |
| | Adult male | 5,159 | 2,030 | 3,129 | 43 | 84 |
| | Larva female | 4,789 | 2,303 | 2,486 | 83 | 414 |
| | Larva male | 5,034 | 2,356 | 2,678 | 67 | 185 |
| *L. reali* | Adult female | 5,039 | 1,947 | 3,092 | 50 | 197 |
| | Adult male | 5,187 | 1,945 | 3,242 | 30 | 69 |
| | Larva female | 4,819 | 2,242 | 2,577 | 97 | 370 |
| | Larva male | 5,081 | 2,261 | 2,820 | 55 | 150 |

**Supplementary table 2**. Gene annotation statistics.

| Species | Predicted genes (N) | Median gene size (bp) | Genes on chromosome scaffolds (%) | Genes with functional predictions (N) |
|---|---|---|---|---|
| *L. juvernica* | 16,417 | 5,868 | 90.87% | 9,816 |
| *L. sinapis* | 16,603 | 6,000 | 88.00% | 9,852 |
| *L. reali* | 15,921 | 6,235 | 85.59% | 9,509 |

**Supplementary table 3.** Number of sex-biased genes in each species, stage and tissue.

| Species | Stage | Tissue | Female-biased | Unbiased | Male-biased |
|---|---|---|---|---|---|
| *L. juvernica* | Larva | Abdomen | 30 | 10,768 | 576 |
| | | Head | 3 | 11,371 | 0 |
| | Adult | Abdomen | 2,144 | 6,940 | 3,374 |
| | | Head | 36 | 12,414 | 8 |
| *L. sinapis* | Larva | Abdomen | 24 | 10,657 | 630 |
| | | Head | 5 | 11,306 | 0 |
| | Adult | Abdomen | 2,200 | 6,720 | 3,556 |
| | | Head | 71 | 12,400 | 5 |
| *L. reali* | Larva | Abdomen | 47 | 10,704 | 722 |
| | | Head | 7 | 11,466 | 0 |
| | Adult | Abdomen | 2,463 | 5,944 | 3,961 |
| | | Head | 43 | 12,320 | 5 |

**Supplementary table 4.** Results of Kruskal-Wallis tests for comparing gene expression levels between different Z chromosome regions and the autosomes. Significant test p-values are highlighted in bold.

| Species | sex | stage | tissue | statistic | p-value |
|---|---|---|---|---|---|
| *L. juvernica* | female | adult | abdomen | 116.67 | **< 2.2e-16** |
| | | | head | 72.417 | **3.216e-14** |

| | | | | | |
|---|---|---|---|---|---|
| | | larva | abdomen | 95.641 | **< 2.2e-16** |
| | | | head | 96.657 | **< 2.2e-16** |
| *L. juvernica* | male | adult | abdomen | 50.844 | **9.308e-10** |
| | | | head | 63.036 | **2.862e-12** |
| | | larva | abdomen | 95.156 | **< 2.2e-16** |
| | | | head | 87.33 | **< 2.2e-16** |
| *L. sinapis* | female | adult | abdomen | 158.67 | **< 2.2e-16** |
| | | | head | 83.79 | **< 2.2e-16** |
| | | larva | abdomen | 83.617 | **< 2.2e-16** |
| | | | head | 90.806 | **< 2.2e-16** |
| *L. sinapis* | male | adult | abdomen | 45.78 | **1.007e-08** |
| | | | head | 73.818 | **1.641e-14** |
| | | larva | abdomen | 73.377 | **2.029e-14** |
| | | | head | 84.254 | **< 2.2e-16** |
| *L. reali* | female | adult | abdomen | 119.55 | **< 2.2e-16** |
| | | | head | 64.145 | **1.686e-12** |
| | | larva | abdomen | 88.183 | **< 2.2e-16** |
| | | | head | 77.127 | **3.346e-15** |
| *L. reali* | male | adult | abdomen | 29.251 | **2.070e-05** |
| | | | head | 62.799 | **3.205e-12** |
| | | larva | abdomen | 76.616 | **4.279e-15** |
| | | | head | 65.944 | **7.138e-13** |

**Supplementary table 5.** Results of pairwise Wilcoxon rank sum tests of difference in gene expression levels between Z-linked genes and autosomal genes for *L. juvernica* **A**, female larva abdomen; **B**, female larva head; **C**, female adult abdomen; **D**, female adult head; **E**, male larva abdomen; **F** male larva head; **G**, male adult abdomen and **H**, male adult head. Significant test p-values are highlighted in bold.

**A**

| group1 | group2 | n1 | n2 | statistic | p-value | p.adj | p.adj.signif |
|---|---|---|---|---|---|---|---|
| A | Z1a | 9,323 | 398 | 2237962 | 2.96e-12 | **1.47e-10** | *** |
| A | Z1b | 9,323 | 184 | 869294 | 0.753 | 1 | |
| A | Z2a | 9,323 | 230 | 1214092 | 5.92e-4 | **0.004** | ** |
| A | Z2b | 9,323 | 229 | 1325668 | 3.79e-10 | **9.43e-09** | *** |
| A | Z3 | 9,323 | 286 | 1383638 | 0.275 | 1 | |
| Z1a | Z1b | 398 | 184 | 29321 | 1.1e-4 | **9.13e-4** | *** |
| Z1a | Z2a | 398 | 230 | 42248 | 0.108 | 0.489 | |
| Z1a | Z2b | 398 | 229 | 47034 | 0.503 | 1 | |
| Z1a | Z3 | 398 | 286 | 46672 | 5.88e-05 | **5.85e-4** | *** |
| Z1b | Z2a | 184 | 230 | 23762 | 0.032 | 0.18 | |
| Z1b | Z2b | 184 | 229 | 26072 | 3.33e-05 | **4.14e-4** | *** |
| Z1b | Z3 | 184 | 286 | 27075 | 0.596 | 1 | |
| Z2a | Z2b | 230 | 229 | 29373 | 0.032 | 0.18 | |
| Z2a | Z3 | 230 | 286 | 29492 | 0.044 | 0.217 | |
| Z2b | Z3 | 229 | 286 | 25229 | 7.48e-06 | **1.24e-4** | *** |

**B**

| group1 | group2 | n1 | n2 | statistic | p-value | p.adj | p.adj.signif |
|---|---|---|---|---|---|---|---|
| A | Z1a | 9,420 | 409 | 2317382 | 3.41e-12 | **1.7e-10** | *** |
| A | Z1b | 9,420 | 187 | 903348 | 0.548 | 1 | |
| A | Z2a | 9,420 | 234 | 1233958 | 0.002 | **0.012** | * |
| A | Z2b | 9,420 | 233 | 1367245 | 1.35e-10 | **3.36e-09** | *** |
| A | Z3 | 9,420 | 292 | 1459194 | 0.076 | 0.376 | |
| Z1a | Z1b | 409 | 187 | 31153 | 2.79e-4 | **0.003** | ** |
| Z1a | Z2a | 409 | 234 | 43416 | 0.05 | 0.278 | |

| | | | | | | | |
|---|---|---|---|---|---|---|---|
| Z1a | Z2b | 409 | 233 | 48744 | 0.628 | 1 | |
| Z1a | Z3 | 409 | 292 | 50663 | 6.17e-4 | **0.005** | ** |
| Z1b | Z2a | 187 | 234 | 24025 | 0.084 | 0.379 | |
| Z1b | Z2b | 187 | 233 | 26788 | 5.22e-05 | **8.66e-4** | *** |
| Z1b | Z3 | 187 | 292 | 28378 | 0.467 | 1 | |
| Z2a | Z2b | 234 | 233 | 30768 | 0.016 | 0.101 | |
| Z2a | Z3 | 234 | 292 | 32011 | 0.214 | 0.888 | |
| Z2b | Z3 | 233 | 292 | 27193 | 7.76e-05 | **9.66e-4** | *** |

**C**

| group1 | group2 | n1 | n2 | statistic | p-value | p.adj | p.adj.signif |
|---|---|---|---|---|---|---|---|
| A | Z1a | 10,195 | 472 | 2956476 | 3.89e-17 | **1.94e-15** | *** |
| A | Z1b | 10,195 | 218 | 1169861 | 0.182 | 0.824 | |
| A | Z2a | 10,195 | 257 | 1420090 | 0.021 | 0.106 | |
| A | Z2b | 10,195 | 269 | 1678317 | 3.4e-10 | **8.46e-09** | *** |
| A | Z3 | 10,195 | 310 | 1728442 | 0.005 | **0.028** | * |
| Z1a | Z1b | 472 | 218 | 42275 | 1.64e-4 | **0.003** | ** |
| Z1a | Z2a | 472 | 257 | 51385 | 6.47e-4 | **0.006** | ** |
| Z1a | Z2b | 472 | 269 | 62208 | 0.649 | 1 | |
| Z1a | Z3 | 472 | 310 | 62603 | 6.34e-4 | **0.006** | ** |
| Z1b | Z2a | 218 | 257 | 28806 | 0.595 | 1 | |
| Z1b | Z2b | 218 | 269 | 34265 | 0.001 | **0.011** | * |
| Z1b | Z3 | 218 | 310 | 35078 | 0.456 | 1 | |
| Z2a | Z2b | 257 | 269 | 39631 | 0.004 | **0.026** | * |
| Z2a | Z3 | 257 | 310 | 40312 | 0.806 | 1 | |
| Z2b | Z3 | 269 | 310 | 36062 | 0.005 | **0.028** | * |

**D**

| group1 | group2 | n1 | n2 | statistic | p-value | p.adj | p.adj.signif |
|---|---|---|---|---|---|---|---|
| A | Z1a | 10,168 | 476 | 2672733 | 1.15e-4 | **0.002** | ** |
| A | Z1b | 10,168 | 221 | 1207659 | 0.057 | 0.352 | |
| A | Z2a | 10,168 | 263 | 1387532 | 0.296 | 1 | |
| A | Z2b | 10,168 | 266 | 1687677 | 4.69e-12 | **2.33e-10** | **** |
| A | Z3 | 10,168 | 307 | 1729145 | 0.001 | **0.012** | * |
| Z1a | Z1b | 476 | 221 | 51151 | 0.559 | 1 | |
| Z1a | Z2a | 476 | 263 | 58371 | 0.129 | 0.713 | |
| Z1a | Z2b | 476 | 266 | 72252 | 0.001 | **0.012** | * |
| Z1a | Z3 | 476 | 307 | 73058 | 0.998 | 1 | |
| Z1b | Z2a | 221 | 263 | 27980 | 0.481 | 1 | |
| Z1b | Z2b | 221 | 266 | 34482 | 9.98e-4 | **0.012** | * |
| Z1b | Z3 | 221 | 307 | 34950 | 0.553 | 1 | |
| Z2a | Z2b | 263 | 266 | 42663 | 1.24e-05 | **3.09e-4** | *** |
| Z2a | Z3 | 263 | 307 | 43191 | 0.15 | 0.747 | |
| Z2b | Z3 | 266 | 307 | 34721 | 0.002 | **0.014** | * |

**E**

| group1 | group2 | n1 | n2 | statistic | p-value | p.adj | p.adj.signif |
|---|---|---|---|---|---|---|---|
| A | Z1a | 9,516 | 416 | 2365761 | 1.47e-11 | **3.66e-10** | *** |
| A | Z1b | 9,516 | 191 | 911019 | 0.953 | 1 | |
| A | Z2a | 9,516 | 240 | 1265537 | 0.004 | **0.026** | * |
| A | Z2b | 9,516 | 233 | 1395410 | 1.41e-11 | **3.66e-10** | *** |
| A | Z3 | 9,516 | 293 | 1400014 | 0.901 | 1 | |
| Z1a | Z1b | 416 | 191 | 31946 | 1.05e-4 | **8.71e-4** | *** |
| Z1a | Z2a | 416 | 240 | 45617 | 0.066 | 0.297 | |
| Z1a | Z2b | 416 | 233 | 52085 | 0.114 | 0.473 | |
| Z1a | Z3 | 416 | 293 | 49109 | 1.05e-05 | **1.05e-4** | *** |
| Z1b | Z2a | 191 | 240 | 25399 | 0.054 | 0.267 | |
| Z1b | Z2b | 191 | 233 | 28082 | 3.42e-06 | **4.26e-05** | *** |
| Z1b | Z3 | 191 | 293 | 28081 | 0.948 | 1 | |
| Z2a | Z2b | 240 | 233 | 32311 | 0.003 | **0.024** | * |

| | | | | | | | |
|---|---|---|---|---|---|---|---|
| Z2a | Z3 | 240 | 293 | 31467 | 0.037 | 0.204 | |
| Z2b | Z3 | 233 | 293 | 25194 | 2.43e-07 | **4.03e-06** | *** |

**F**

| group1 | group2 | n1 | n2 | statistic | p-value | p.adj | p.adj.signif |
|---|---|---|---|---|---|---|---|
| A | Z1a | 9,384 | 407 | 2291010 | 8.41e-12 | **4.19e-10** | *** |
| A | Z1b | 9,384 | 188 | 899933 | 0.634 | 1 | |
| A | Z2a | 9,384 | 228 | 1163350 | 0.024 | 0.132 | |
| A | Z2b | 9,384 | 230 | 1340182 | 3.46e-10 | **8.61e-09** | *** |
| A | Z3 | 9,384 | 291 | 1375910 | 0.822 | 1 | |
| Z1a | Z1b | 407 | 188 | 31223 | 3.08e-4 | **0.003** | ** |
| Z1a | Z2a | 407 | 228 | 40963 | 0.014 | 0.089 | |
| Z1a | Z2b | 407 | 230 | 48401 | 0.474 | 1 | |
| Z1a | Z3 | 407 | 291 | 47639 | 1.04e-05 | **1.29e-4** | *** |
| Z1b | Z2a | 188 | 228 | 22861 | 0.242 | 1 | |
| Z1b | Z2b | 188 | 230 | 26447 | 8.57e-05 | **8.53e-4** | *** |
| Z1b | Z3 | 188 | 291 | 27126 | 0.878 | 1 | |
| Z2a | Z2b | 228 | 230 | 30474 | 0.003 | **0.019** | * |
| Z2a | Z3 | 228 | 291 | 30478 | 0.112 | 0.557 | |
| Z2b | Z3 | 230 | 291 | 25463 | 2.74e-06 | **4.55e-05** | *** |

**G**

| group1 | group2 | n1 | n2 | statistic | p-value | p.adj | p.adj.signif |
|---|---|---|---|---|---|---|---|
| A | Z1a | 10,308 | 490 | 2806323 | 3.1e-05 | **5.14e-4** | *** |
| A | Z1b | 10,308 | 227 | 1091723 | 0.084 | 0.381 | |
| A | Z2a | 10,308 | 271 | 1389137 | 0.878 | 1 | |
| A | Z2b | 10,308 | 273 | 1681562 | 3.57e-08 | **1.78e-06** | *** |
| A | Z3 | 10,308 | 309 | 1643188 | 0.341 | 1 | |
| Z1a | Z1b | 490 | 227 | 45848 | 1.53e-4 | **0.002** | ** |
| Z1a | Z2a | 490 | 271 | 58800 | 0.009 | 0.063 | |
| Z1a | Z2b | 490 | 273 | 72706 | 0.046 | 0.255 | |
| Z1a | Z3 | 490 | 309 | 69303 | 0.044 | 0.255 | |
| Z1b | Z2a | 227 | 271 | 32654 | 0.236 | 0.979 | |
| Z1b | Z2b | 227 | 273 | 39043 | 5.47e-07 | **1.36e-05** | *** |
| Z1b | Z3 | 227 | 309 | 38447 | 0.057 | 0.283 | |
| Z2a | Z2b | 271 | 273 | 44338 | 6.14e-05 | **7.64e-4** | *** |
| Z2a | Z3 | 271 | 309 | 43328 | 0.469 | 1 | |
| Z2b | Z3 | 273 | 309 | 34720 | 2.3e-4 | **0.002** | ** |

**H**

| group1 | group2 | n1 | n2 | statistic | p-value | p.adj | p.adj.signif |
|---|---|---|---|---|---|---|---|
| A | Z1a | 10,149 | 481 | 2728268 | 1.24e-05 | **3.09e-4** | *** |
| A | Z1b | 10,149 | 225 | 1200572 | 0.186 | 0.931 | |
| A | Z2a | 10,149 | 263 | 1369843 | 0.464 | 1 | |
| A | Z2b | 10,149 | 269 | 1677825 | 1.32e-10 | **6.57e-09** | *** |
| A | Z3 | 10,149 | 298 | 1621959 | 0.032 | 0.23 | |
| Z1a | Z1b | 481 | 225 | 50779 | 0.187 | 0.931 | |
| Z1a | Z2a | 481 | 263 | 57573 | 0.043 | 0.266 | |
| Z1a | Z2b | 481 | 269 | 71647 | 0.015 | 0.121 | |
| Z1a | Z3 | 481 | 298 | 67904 | 0.217 | 0.982 | |
| Z1b | Z2a | 225 | 263 | 28826 | 0.624 | 1 | |
| Z1b | Z2b | 225 | 269 | 35509 | 9,00e-04 | **0.009** | ** |
| Z1b | Z3 | 225 | 298 | 34042 | 0.763 | 1 | |
| Z2a | Z2b | 263 | 269 | 42555 | 5.1e-05 | **8.46e-4** | *** |
| Z2a | Z3 | 263 | 298 | 40911 | 0.368 | 1 | |
| Z2b | Z3 | 269 | 298 | 33105 | 3.42e-4 | **0.004** | ** |

Note – The different Z chromosomes are divided into a and b regions according to previously inferred stepwise fusions of autosomal parts to the Z chromosomes. Correction for multiple testing was done with Benjamini-Yekutieli correction. Significant adjusted p-values are highlighted in bold and asterisks denote significance levels (* $P \le 0.05$, ** $P \le 0.01$, *** $P \le 0.001$).

**Supplementary table 6.** Results of pairwise Wilcoxon rank sum tests of difference in gene expression levels between Z-linked genes and autosomal genes for *L. sinapis* **A**, female larva abdomen; **B**, female larva head; **C**, female adult abdomen; **D**, female adult head; **E**, male larva abdomen; **F,** male larva head; **G**, male adult abdomen and **H**, male adult head. Significant test p-values are highlighted in bold.

**A**

| group1 | group2 | n1 | n2 | statistic | p-value | p.adj | p.adj.signif |
|---|---|---|---|---|---|---|---|
| A | Z1a | 8,954 | 417 | 2289101 | 5.36e-15 | **2.67e-13** | *** |
| A | Z1b | 8,954 | 186 | 856289 | 0.508 | 1 | |
| A | Z2a | 8,954 | 199 | 984147 | 0.012 | 0.072 | |
| A | Z2b | 8,954 | 211 | 1110335 | 1.29e-05 | **2.14e-04** | *** |
| A | Z3 | 8,954 | 258 | 1150666 | 0.917 | 1 | |
| Z1a | Z1b | 417 | 186 | 30876 | 6.32e-05 | **7.86e-04** | *** |
| Z1a | Z2a | 417 | 199 | 36050 | 0.008 | 0.07 | |
| Z1a | Z2b | 417 | 211 | 40911 | 0.151 | 0.752 | |
| Z1a | Z3 | 417 | 258 | 41095 | 2.5e-07 | **6.22e-06** | *** |
| Z1b | Z2a | 186 | 199 | 19902 | 0.201 | 0.834 | |
| Z1b | Z2b | 186 | 211 | 22526 | 0.011 | 0.072 | |
| Z1b | Z3 | 186 | 258 | 23178 | 0.541 | 1 | |
| Z2a | Z2b | 199 | 211 | 22606 | 0.179 | 0.81 | |
| Z2a | Z3 | 199 | 258 | 22731 | 0.036 | 0.197 | |
| Z2b | Z3 | 211 | 258 | 21999 | 0.000351 | **0.003** | ** |

**B**

| group1 | group2 | n1 | n2 | statistic | p-value | p.adj | p.adj.signif |
|---|---|---|---|---|---|---|---|
| A | Z1a | 9,044 | 422 | 2343583 | 2.14e-15 | **1.07e-13** | *** |
| A | Z1b | 9,044 | 191 | 937271 | 0.044 | 0.271 | |
| A | Z2a | 9,044 | 207 | 1031686 | 0.012 | 0.084 | |
| A | Z2b | 9,044 | 214 | 1150868 | 2.14e-06 | **3.55e-05** | *** |
| A | Z3 | 9,044 | 258 | 1177525 | 0.799 | 1 | |
| Z1a | Z1b | 422 | 191 | 34369 | 0.003 | **0.035** | * |
| Z1a | Z2a | 422 | 207 | 37976 | 0.008 | 0.064 | |
| Z1a | Z2b | 422 | 214 | 42286 | 0.19 | 0.727 | |
| Z1a | Z3 | 422 | 258 | 42011 | 5.76e-07 | **1.43e-05** | *** |
| Z1b | Z2a | 191 | 207 | 20097 | 0.775 | 1 | |
| Z1b | Z2b | 191 | 214 | 22505 | 0.079 | 0.392 | |
| Z1b | Z3 | 191 | 258 | 22642 | 0.142 | 0.643 | |
| Z2a | Z2b | 207 | 214 | 23883 | 0.165 | 0.684 | |
| Z2a | Z3 | 207 | 258 | 24039 | 0.064 | 0.356 | |
| Z2b | Z3 | 214 | 258 | 22449 | 0.000473 | **0.006** | ** |

**C**

| group1 | group2 | n1 | n2 | statistic | p-value | p.adj | p.adj.signif |
|---|---|---|---|---|---|---|---|
| A | Z1a | 9,831 | 502 | 3199819 | 2.84e-29 | **1.41e-27** | *** |
| A | Z1b | 9,831 | 214 | 1094541 | 0.31 | 1 | |
| A | Z2a | 9,831 | 233 | 1269732 | 0.005 | **0.032** | * |
| A | Z2b | 9,831 | 256 | 1493709 | 3.11e-07 | **3.87e-06** | *** |
| A | Z3 | 9,831 | 288 | 1538505 | 0.012 | 0.066 | |
| Z1a | Z1b | 502 | 214 | 39585 | 2.45e-08 | **6.10e-07** | *** |
| Z1a | Z2a | 502 | 233 | 46941 | 1.64e-05 | **1.63e-04** | *** |
| Z1a | Z2b | 502 | 256 | 54924 | 0.001 | **0.009** | ** |
| Z1a | Z3 | 502 | 288 | 55859 | 1.03e-07 | **1.71e-06** | *** |
| Z1b | Z2a | 214 | 233 | 26665 | 0.204 | 0.846 | |
| Z1b | Z2b | 214 | 256 | 31354 | 0.007 | **0.043** | * |
| Z1b | Z3 | 214 | 288 | 32188 | 0.393 | 1 | |
| Z2a | Z2b | 233 | 256 | 32083 | 0.148 | 0.67 | |
| Z2a | Z3 | 233 | 288 | 32813 | 0.666 | 1 | |

| | | | | | | |
|---|---|---|---|---|---|---|
| Z2b | Z3 | 256 | 288 | 33229 | 0.047 | 0.234 |

**D**

| group1 | group2 | n1 | n2 | statistic | p-value | p.adj | p.adj.signif |
|---|---|---|---|---|---|---|---|
| A | Z1a | 9,786 | 491 | 2769153 | 1.09e-08 | **2.71e-07** | *** |
| A | Z1b | 9,786 | 215 | 1097663 | 0.275 | 1 | |
| A | Z2a | 9,786 | 235 | 1194207 | 0.312 | 1 | |
| A | Z2b | 9,786 | 257 | 1568530 | 1.21e-11 | **6.02e-10** | *** |
| A | Z3 | 9,786 | 286 | 1542730 | 0.003 | **0.026** | * |
| Z1a | Z1b | 491 | 215 | 46855 | 0.018 | 0.109 | |
| Z1a | Z2a | 491 | 235 | 50584 | 0.007 | 0.051 | |
| Z1a | Z2b | 491 | 257 | 68938 | 0.037 | 0.206 | |
| Z1a | Z3 | 491 | 286 | 66336 | 0.199 | 0.9 | |
| Z1b | Z2a | 215 | 235 | 25068 | 0.888 | 1 | |
| Z1b | Z2b | 215 | 257 | 33320 | 0.000115 | **0.001** | ** |
| Z1b | Z3 | 215 | 286 | 32465 | 0.284 | 1 | |
| Z2a | Z2b | 235 | 257 | 36978 | 1.68e-05 | **2.79e-04** | *** |
| Z2a | Z3 | 235 | 286 | 35808 | 0.198 | 0.9 | |
| Z2b | Z3 | 257 | 286 | 30961 | 0.002 | **0.015** | * |

**E**

| group1 | group2 | n1 | n2 | statistic | p-value | p.adj | p.adj.signif |
|---|---|---|---|---|---|---|---|
| A | Z1a | 9,169 | 438 | 2386627 | 2.44e-11 | **1.21e-09** | *** |
| A | Z1b | 9,169 | 195 | 920100 | 0.484 | 1 | |
| A | Z2a | 9,169 | 211 | 1073432 | 0.006 | **0.045** | * |
| A | Z2b | 9,169 | 216 | 1180344 | 1.37e-06 | **2.27e-05** | *** |
| A | Z3 | 9,169 | 263 | 1170085 | 0.413 | 1 | |
| Z1a | Z1b | 438 | 195 | 35989 | 0.002 | **0.016** | * |
| Z1a | Z2a | 438 | 211 | 42470 | 0.095 | 0.524 | |
| Z1a | Z2b | 438 | 216 | 47474 | 0.941 | 1 | |
| Z1a | Z3 | 438 | 263 | 44969 | 1.15e-06 | **2.27e-05** | *** |
| Z1b | Z2a | 195 | 211 | 22279 | 0.149 | 0.674 | |
| Z1b | Z2b | 195 | 216 | 24412 | 0.005 | **0.044** | * |
| Z1b | Z3 | 195 | 263 | 24184 | 0.298 | 1 | |
| Z2a | Z2b | 211 | 216 | 24717 | 0.13 | 0.647 | |
| Z2a | Z3 | 211 | 263 | 23900 | 0.009 | 0.059 | |
| Z2b | Z3 | 216 | 263 | 22081 | 2.74e-05 | **3.41e-04** | *** |

**F**

| group1 | group2 | n1 | n2 | statistic | p-value | p.adj | p.adj.signif |
|---|---|---|---|---|---|---|---|
| A | Z1a | 9,057 | 429 | 2379367 | 3.32e-15 | **1.65e-13** | *** |
| A | Z1b | 9,057 | 189 | 903450 | 0.19 | 0.747 | |
| A | Z2a | 9,057 | 206 | 1008877 | 0.045 | 0.281 | |
| A | Z2b | 9,057 | 216 | 1150841 | 8.95e-06 | **1.48e-04** | *** |
| A | Z3 | 9,057 | 258 | 1150990 | 0.684 | 1 | |
| Z1a | Z1b | 429 | 189 | 33445 | 0.000522 | **0.005** | ** |
| Z1a | Z2a | 429 | 206 | 37355 | 0.002 | **0.013** | * |
| Z1a | Z2b | 429 | 216 | 43154 | 0.155 | 0.701 | |
| Z1a | Z3 | 429 | 258 | 41950 | 1.06e-07 | **2.64e-06** | *** |
| Z1b | Z2a | 189 | 206 | 19987 | 0.647 | 1 | |
| Z1b | Z2b | 189 | 216 | 22889 | 0.035 | 0.25 | |
| Z1b | Z3 | 189 | 258 | 22632 | 0.195 | 0.747 | |
| Z2a | Z2b | 206 | 216 | 24466 | 0.077 | 0.387 | |
| Z2a | Z3 | 206 | 258 | 24042 | 0.078 | 0.387 | |
| Z2b | Z3 | 216 | 258 | 22401 | 0.000235 | **0.003** | ** |

**G**

| group1 | group2 | n1 | n2 | statistic | p-value | p.adj | p.adj.signif |
|---|---|---|---|---|---|---|---|
| A | Z1a | 10,003 | 510 | 2885958 | 5.34e-07 | **2.66e-05** | *** |

| | | | | | | | |
|---|---|---|---|---|---|---|---|
| A | Z1b | 10,003 | 221 | 1024290 | 0.062 | 0.308 | |
| A | Z2a | 10,003 | 248 | 1268215 | 0.545 | 1 | |
| A | Z2b | 10,003 | 268 | 1537887 | 3.75e-05 | **6.22e-04** | *** |
| A | Z3 | 10,003 | 292 | 1511930 | 0.304 | 1 | |
| Z1a | Z1b | 510 | 221 | 45211 | 2.14e-05 | **5.33e-04** | *** |
| Z1a | Z2a | 510 | 248 | 56463 | 0.017 | 0.133 | |
| Z1a | Z2b | 510 | 268 | 69186 | 0.777 | 1 | |
| Z1a | Z3 | 510 | 292 | 67190 | 0.021 | 0.133 | |
| Z1b | Z2a | 221 | 248 | 29951 | 0.082 | 0.372 | |
| Z1b | Z2b | 221 | 268 | 35821 | 6.58e-05 | **8.19e-04** | *** |
| Z1b | Z3 | 221 | 292 | 35638 | 0.043 | 0.236 | |
| Z2a | Z2b | 248 | 268 | 37354 | 0.015 | 0.133 | |
| Z2a | Z3 | 248 | 292 | 36635 | 0.813 | 1 | |
| Z2b | Z3 | 268 | 292 | 34716 | 0.021 | 0.133 | |

**H**

| group1 | group2 | n1 | n2 | statistic | p-value | p.adj | p.adj.signif |
|---|---|---|---|---|---|---|---|
| A | Z1a | 9,791 | 497 | 2810151 | 5.29e-09 | **1.32e-07** | *** |
| A | Z1b | 9,791 | 215 | 1084817 | 0.441 | 1 | |
| A | Z2a | 9,791 | 238 | 1225340 | 0.172 | 0.778 | |
| A | Z2b | 9,791 | 258 | 1548448 | 5.47e-10 | **2.72e-08** | *** |
| A | Z3 | 9,791 | 283 | 1478321 | 0.054 | 0.299 | |
| Z1a | Z1b | 497 | 215 | 46596 | 0.007 | 0.056 | |
| Z1a | Z2a | 497 | 238 | 52884 | 0.02 | 0.143 | |
| Z1a | Z2b | 497 | 258 | 68141 | 0.156 | 0.776 | |
| Z1a | Z3 | 497 | 283 | 63790 | 0.031 | 0.192 | |
| Z1b | Z2a | 215 | 238 | 26132 | 0.694 | 1 | |
| Z1b | Z2b | 215 | 258 | 33290 | 0.000175 | **0.003** | ** |
| Z1b | Z3 | 215 | 283 | 31464 | 0.513 | 1 | |
| Z2a | Z2b | 238 | 258 | 36066 | 0.00077 | **0.008** | ** |
| Z2a | Z3 | 238 | 283 | 34019 | 0.842 | 1 | |
| Z2b | Z3 | 258 | 283 | 30275 | 6e-04 | **0.007** | ** |

Note – The different Z chromosomes are divided into a and b regions according to previously inferred stepwise fusions of autosomal parts to the Z chromosomes. Correction for multiple testing was done with Benjamini-Yekutieli correction. Significant adjusted p-values are highlighted in bold and asterisks denote significance levels (* $P \leq 0.05$, ** $P \leq 0.01$, *** $P \leq 0.001$).

**Supplementary table 7.** Results of pairwise Wilcoxon rank sum tests of difference in gene expression levels between Z-linked genes and autosomal genes for *L. reali* **A**, female larva abdomen; **B**, female larva head; **C**, female adult abdomen; **D**, female adult head; **E**, male larva abdomen; **F,** male larva head; **G**, male adult abdomen and **H**, male adult head. Significant test p-values are highlighted in bold.

**A**

| group1 | group2 | n1 | n2 | statistic | p-value | p.adj | p.adj.signif |
|---|---|---|---|---|---|---|---|
| A | Z1a | 8,734 | 412 | 2191341 | 7.02e-14 | **3.49e-12** | *** |
| A | Z1b | 8,734 | 186 | 822442 | 0.77 | 1 | |
| A | Z2a | 8,734 | 206 | 1023366 | 7.24e-4 | **0.007** | ** |
| A | Z2b | 8,734 | 251 | 1293550 | 1.1e-06 | **2.74e-05** | *** |
| A | Z3 | 8,734 | 259 | 1200600 | 0.091 | 0.454 | |
| Z1a | Z1b | 412 | 186 | 30666 | 9.19e-05 | **0.002** | ** |
| Z1a | Z2a | 412 | 206 | 38747 | 0.078 | 0.431 | |
| Z1a | Z2b | 412 | 251 | 49605 | 0.38 | 1 | |
| Z1a | Z3 | 412 | 259 | 44323 | 2.21e-4 | **0.003** | ** |
| Z1b | Z2a | 186 | 206 | 21487 | 0.038 | 0.234 | |
| Z1b | Z2b | 186 | 251 | 27128 | 0.004 | **0.031** | * |
| Z1b | Z3 | 186 | 259 | 25227 | 0.394 | 1 | |
| Z2a | Z2b | 206 | 251 | 27110 | 0.371 | 1 | |
| Z2a | Z3 | 206 | 259 | 24449 | 0.122 | 0.552 | |
| Z2b | Z3 | 251 | 259 | 28109 | 0.008 | 0.059 | |

**B**

| group1 | group2 | n1 | n2 | statistic | p-value | p.adj | p.adj.signif |
|---|---|---|---|---|---|---|---|
| A | Z1a | 8,812 | 426 | 2248985 | 4.51e-12 | **2.24e-10** | *** |
| A | Z1b | 8,812 | 187 | 877825 | 0.125 | 0.741 | |
| A | Z2a | 8,812 | 213 | 1052465 | 0.002 | **0.04** | * |
| A | Z2b | 8,812 | 252 | 1274530 | 6.09e-05 | **0.002** | ** |
| A | Z3 | 8,812 | 264 | 1286196 | 0.003 | **0.042** | * |
| Z1a | Z1b | 426 | 187 | 34301 | 0.006 | 0.061 | |
| Z1a | Z2a | 426 | 213 | 41320 | 0.066 | 0.467 | |
| Z1a | Z2b | 426 | 252 | 49983 | 0.134 | 0.741 | |
| Z1a | Z3 | 426 | 264 | 51055 | 0.042 | 0.348 | |
| Z1b | Z2a | 187 | 213 | 21064 | 0.32 | 1 | |
| Z1b | Z2b | 187 | 252 | 25377 | 0.167 | 0.831 | |
| Z1b | Z3 | 187 | 264 | 25792 | 0.417 | 1 | |
| Z2a | Z2b | 213 | 252 | 27340 | 0.728 | 1 | |
| Z2a | Z3 | 213 | 264 | 27808 | 0.837 | 1 | |
| Z2b | Z3 | 252 | 264 | 32243 | 0.547 | 1 | |

**C**

| group1 | group2 | n1 | n2 | statistic | p-value | p.adj | p.adj.signif |
|---|---|---|---|---|---|---|---|
| A | Z1a | 9,449 | 477 | 2818735 | 2.14e-20 | **1.07e-18** | *** |
| A | Z1b | 9,449 | 209 | 1014683 | 0.494 | 1 | |
| A | Z2a | 9,449 | 232 | 1244550 | 4.15e-4 | **0.004** | ** |
| A | Z2b | 9,449 | 280 | 1510011.5 | 5.33e-05 | **8.84e-4** | *** |
| A | Z3 | 9,449 | 288 | 1534461 | 2.17e-4 | **0.003** | ** |
| Z1a | Z1b | 477 | 209 | 38533 | 2.19e-06 | **5.45e-05** | *** |
| Z1a | Z2a | 477 | 232 | 48508 | 0.008 | **0.048** | * |
| Z1a | Z2b | 477 | 280 | 57913 | 0.002 | **0.018** | * |
| Z1a | Z3 | 477 | 288 | 59762 | 0.003 | **0.018** | * |
| Z1b | Z2a | 209 | 232 | 26884 | 0.048 | 0.24 | |
| Z1b | Z2b | 209 | 280 | 32420 | 0.041 | 0.227 | |
| Z1b | Z3 | 209 | 288 | 33095 | 0.058 | 0.262 | |
| Z2a | Z2b | 232 | 280 | 32445 | 0.983 | 1 | |
| Z2a | Z3 | 232 | 288 | 33141 | 0.876 | 1 | |
| Z2b | Z3 | 280 | 288 | 39998 | 0.869 | 1 | |

**D**

| group1 | group2 | n1 | n2 | statistic | p-value | p.adj | p.adj.signif |
|---|---|---|---|---|---|---|---|
| A | Z1a | 9,448 | 482 | 2533387 | 2.96e-05 | **7.37e-4** | *** |
| A | Z1b | 9,448 | 213 | 1056423 | 0.212 | 0.879 | |
| A | Z2a | 9,448 | 234 | 1178608 | 0.083 | 0.591 | |
| A | Z2b | 9,448 | 282 | 1602114 | 6.34e-09 | **3.16e-07** | *** |
| A | Z3 | 9,448 | 283 | 1512926 | 1.57e-4 | **0.003** | ** |
| Z1a | Z1b | 482 | 213 | 48068 | 0.181 | 0.879 | |
| Z1a | Z2a | 482 | 234 | 53457 | 0.258 | 0.988 | |
| Z1a | Z2b | 482 | 282 | 73696 | 0.051 | 0.426 | |
| Z1a | Z3 | 482 | 283 | 69328 | 0.703 | 1 | |
| Z1b | Z2a | 213 | 234 | 25270 | 0.798 | 1 | |
| Z1b | Z2b | 213 | 282 | 34627 | 0.004 | **0.035** | * |
| Z1b | Z3 | 213 | 283 | 32612 | 0.118 | 0.68 | |
| Z2a | Z2b | 234 | 282 | 37973 | 0.003 | **0.035** | * |
| Z2a | Z3 | 234 | 283 | 35289 | 0.198 | 0.879 | |
| Z2b | Z3 | 282 | 283 | 36908 | 0.123 | 0.68 | |

**E**

| group1 | group2 | n1 | n2 | statistic | p-value | p.adj | p.adj.signif |
|---|---|---|---|---|---|---|---|
| A | Z1a | 8,977 | 443 | 2363782 | 1.84e-11 | **9.16e-10** | *** |
| A | Z1b | 8,977 | 194 | 872260 | 0.967 | 1 | |
| A | Z2a | 8,977 | 218 | 1100207 | 0.002 | **0.014** | * |

| | | | | | | | |
|---|---|---|---|---|---|---|---|
| A | Z2b | 8,977 | 254 | 1350489 | 5.07e-07 | **1.26e-05** | *** |
| A | Z3 | 8,977 | 267 | 1248633 | 0.243 | 1 | |
| Z1a | Z1b | 443 | 194 | 35126 | 2.43e-4 | **0.004** | ** |
| Z1a | Z2a | 443 | 218 | 45470 | 0.222 | 1 | |
| Z1a | Z2b | 443 | 254 | 56274 | 0.996 | 1 | |
| Z1a | Z3 | 443 | 267 | 49971 | 5.33e-4 | **0.007** | ** |
| Z1b | Z2a | 194 | 218 | 23723 | 0.033 | 0.203 | |
| Z1b | Z2b | 194 | 254 | 29094 | 0.001 | **0.01** | * |
| Z1b | Z3 | 194 | 267 | 26961 | 0.452 | 1 | |
| Z2a | Z2b | 218 | 254 | 29163 | 0.318 | 1 | |
| Z2a | Z3 | 218 | 267 | 26529 | 0.094 | 0.518 | |
| Z2b | Z3 | 254 | 267 | 28718 | 0.003 | **0.018** | * |

**F**

| group1 | group2 | n1 | n2 | statistic | p-value | p.adj | p.adj.signif |
|---|---|---|---|---|---|---|---|
| A | Z1a | 8,842 | 426 | 2236296 | 6.01e-11 | **2.99e-09** | **** |
| A | Z1b | 8,842 | 187 | 844169 | 0.621 | 1 | |
| A | Z2a | 8,842 | 218 | 1073950 | 0.004 | **0.039** | * |
| A | Z2b | 8,842 | 253 | 1290160 | 3.07e-05 | **7.64e-4** | *** |
| A | Z3 | 8,842 | 260 | 1205955 | 0.176 | 0.876 | |
| Z1a | Z1b | 426 | 187 | 32963 | 6.7e-4 | **0.011** | * |
| Z1a | Z2a | 426 | 218 | 42817 | 0.106 | 0.586 | |
| Z1a | Z2b | 426 | 253 | 51396 | 0.313 | 1 | |
| Z1a | Z3 | 426 | 260 | 47941 | 0.003 | **0.039** | * |
| Z1b | Z2a | 187 | 218 | 22441 | 0.08 | 0.496 | |
| Z1b | Z2b | 187 | 253 | 27056 | 0.01 | 0.082 | |
| Z1b | Z3 | 187 | 260 | 25064 | 0.576 | 1 | |
| Z2a | Z2b | 218 | 253 | 28526 | 0.52 | 1 | |
| Z2a | Z3 | 218 | 260 | 26567 | 0.239 | 1 | |
| Z2b | Z3 | 253 | 260 | 29693 | 0.057 | 0.405 | |

**G**

| group1 | group2 | n1 | n2 | statistic | p-value | p.adj | p.adj.signif |
|---|---|---|---|---|---|---|---|
| A | Z1a | 9,618 | 498 | 2615231 | 5.25e-4 | **0.013** | * |
| A | Z1b | 9,618 | 222 | 1023259 | 0.289 | 1 | |
| A | Z2a | 9,618 | 243 | 1209227 | 0.354 | 1 | |
| A | Z2b | 9,618 | 294 | 1599935 | 1.18e-4 | **0.006** | ** |
| A | Z3 | 9,618 | 290 | 1460249 | 0.171 | 0.946 | |
| Z1a | Z1b | 498 | 222 | 48121 | 0.005 | 0.068 | |
| Z1a | Z2a | 498 | 243 | 57279 | 0.238 | 1 | |
| Z1a | Z2b | 498 | 294 | 76340 | 0.314 | 1 | |
| Z1a | Z3 | 498 | 290 | 68730 | 0.259 | 1 | |
| Z1b | Z2a | 222 | 243 | 29003 | 0.161 | 0.946 | |
| Z1b | Z2b | 222 | 294 | 38044 | 0.001 | **0.021** | * |
| Z1b | Z3 | 222 | 290 | 34913 | 0.101 | 0.718 | |
| Z2a | Z2b | 243 | 294 | 39024 | 0.065 | 0.539 | |
| Z2a | Z3 | 243 | 290 | 35590 | 0.841 | 1 | |
| Z2b | Z3 | 294 | 290 | 38805 | 0.061 | 0.539 | |

**H**

| group1 | group2 | n1 | n2 | statistic | p-value | p.adj | p.adj.signif |
|---|---|---|---|---|---|---|---|
| A | Z1a | 9,463 | 487 | 2573457 | 1.33e-05 | **3.31e-4** | *** |
| A | Z1b | 9,463 | 214 | 1044823 | 0.424 | 1 | |
| A | Z2a | 9,463 | 236 | 1199689 | 0.051 | 0.317 | |
| A | Z2b | 9,463 | 290 | 1653531 | 2.55e-09 | **1.27e-07** | *** |
| A | Z3 | 9,463 | 280 | 1466754 | 0.002 | **0.027** | * |
| Z1a | Z1b | 487 | 214 | 47751 | 0.078 | 0.429 | |

| | | | | | | | |
|---|---|---|---|---|---|---|---|
| Z1a | Z2a | 487 | 236 | 54817 | 0.315 | 1 | |
| Z1a | Z2b | 487 | 290 | 76521 | 0.051 | 0.317 | |
| Z1a | Z3 | 487 | 280 | 67251 | 0.753 | 1 | |
| Z1b | Z2a | 214 | 236 | 26334 | 0.432 | 1 | |
| Z1b | Z2b | 214 | 290 | 36361 | 9.72e-4 | **0.016** | * |
| Z1b | Z3 | 214 | 280 | 32226 | 0.15 | 0.747 | |
| Z2a | Z2b | 236 | 290 | 38901 | 0.007 | 0.069 | |
| Z2a | Z3 | 236 | 280 | 34050 | 0.55 | 1 | |
| Z2b | Z3 | 290 | 280 | 36273 | 0.028 | 0.23 | |

Note – The different Z chromosomes are divided into a and b regions according to previously inferred stepwise fusions of autosomal parts to the Z chromosomes. Correction for multiple testing was done with Benjamini-Yekutieli correction. Significant adjusted p-values are highlighted in bold and asterisks denote significance levels (* $P \leq 0.05$, ** $P \leq 0.01$, *** $P \leq 0.001$).

**Supplementary table 8.** *L. juvernica* Z:A ratios of median gene expression levels (TPM) for different Z-linked regions.

| | Female | | Male | |
|---|---|---|---|---|
| | **Abdomen** | **Head** | **Abdomen** | **Head** |
| **Larva** | | | | |
| Z1a | 0.47 (0.39-0.58) | 0.59 (0.47-0.69) | 0.58 (0.48-0.68) | 0.58 (0.46-0.71) |
| Z1b | 1.02 (0.68-1.40) | 0.93 (0.73-1.33) | 0.97 (0.76-1.27) | 0.99 (0.73-1.41) |
| Z2a | 0.61 (0.43-0.82) | 0.69 (0.57-0.90) | 0.71 (0.56-0.93) | 0.75 (0.59-0.94) |
| Z2b | 0.38 (0.29-0.48) | 0.41 (0.35-0.56) | 0.42 (0.32-0.52) | 0.40 (0.35-0.55) |
| Z3 | 0.79 (0.62-1.05) | 0.79 (0.71-0.97) | 0.96 (0.83-1.21) | 1.04 (0.81-1.20) |
| **Adult** | | | | |
| Z1a | 0.44 (0.35-0.53) | 0.77 (0.67-0.89) | 0.71 (0.55-0.85) | 0.73 (0.62-0.88) |
| Z1b | 0.81 (0.56-1.12) | 0.79 (0.57-1.01) | 1.09 (0.86-1.37) | 0.74 (0.64-1.07) |
| Z2a | 0.77 (0.58-1.06) | 0.90 (0.73-1.03) | 0.94 (0.74-1.24) | 0.86 (0.71-1.06) |
| Z2b | 0.48 (0.36-0.58) | 0.49 (0.43-0.57) | 0.48 (0.37-0.56) | 0.53 (0.42-0.61) |
| Z3 | 0.82 (0.57-0.97) | 0.79 (0.71-0.91) | 0.97 (0.78-1.10) | 0.86 (0.72-1.00) |

Note – The different Z chromosomes are grouped according to previously inferred stepwise fusions of autosomal parts to the Z chromosomes. Confidence intervals in brackets were obtained with stratified bootstrap.

**Supplementary table 9.** *L. sinapis* Z:A ratios of median gene expression levels (TPM) for different Z-linked regions.

| | Female | | Male | |
|---|---|---|---|---|
| | **Abdomen** | **Head** | **Abdomen** | **Head** |
| **Larva** | | | | |
| Z1a | 0.48 (0.36-0.61) | 0.52 (0.41-0.58) | 0.53 (0.42-0.67) | 0.51 (0.42-0.57) |
| Z1b | 0.90 (0.63-1.25) | 0.77 (0.62-1.00) | 0.88 (0.70-1.48) | 0.84 (0.65-1.10) |
| Z2a | 0.65 (0.51-0.84) | 0.72 (0.54-1.07) | 0.65 (0.53-0.95) | 0.79 (0.61-1.00) |
| Z2b | 0.43 (0.32-0.66) | 0.52 (0.45-0.63) | 0.58 (0.38-0.77) | 0.61 (0.42-0.76) |
| Z3 | 0.99 (0.78-1.24) | 0.94 (0.75-1.19) | 1.19 (0.90-1.36) | 1.14 (0.82-1.41) |
| **Adult** | | | | |

| | | | | |
|---|---|---|---|---|
| Z1a | 0.32 (0.25-0.41) | 0.65 (0.54-0.77) | 0.64 (0.51-0.76) | 0.64 (0.55-0.75) |
| Z1b | 0.78 (0.60-1.03) | 0.90 (0.70-1.11) | 1.25 (0.97-1.68) | 0.96 (0.72-1.18) |
| Z2a | 0.69 (0.50-0.89) | 0.88 (0.69-1.10) | 0.86 (0.71-1.13) | 0.85 (0.71-1.04) |
| Z2b | 0.56 (0.46-0.65) | 0.52 (0.41-0.61) | 0.58 (0.44-0.74) | 0.58 (0.46-0.69) |
| Z3 | 0.66 (0.56-0.93) | 0.78 (0.64-0.94) | 0.93 (0.72-1.12) | 0.92 (0.78-1.09) |

Note – The different Z chromosomes are grouped according to previously inferred stepwise fusions of autosomal parts to the Z chromosomes. Confidence intervals in brackets were obtained with stratified bootstrap.

**Supplementary table 10.** *L. reali* Z:A ratios of median gene expression levels (TPM) for different Z-linked regions.

| | **Female** | | **Male** | |
|---|---|---|---|---|
| | **Abdomen** | **Head** | **Abdomen** | **Head** |
| **Larva** | | | | |
| Z1a | 0.38 (0.30-0.57) | 0.57 (0.48-0.71) | 0.53 (0.45-0.63) | 0.63 (0.51-0.72) |
| Z1b | 0.91 (0.62-1.53) | 0.77 (0.63-1.03) | 1.11 (0.74-1.44) | 0.85 (0.69-1.12) |
| Z2a | 0.56 (0.41-0.80) | 0.71 (0.49-0.97) | 0.63 (0.53-0.87) | 0.70 (0.49-0.96) |
| Z2b | 0.42 (0.34-0.59) | 0.66 (0.55-0.79) | 0.55 (0.42-0.68) | 0.60 (0.46-0.75) |
| Z3 | 0.77 (0.52-1.12) | 0.73 (0.55-0.99) | 0.89 (0.71-1.10) | 0.83 (0.61-1.09) |
| **Adult** | | | | |
| Z1a | 0.37 (0.28-0.46) | 0.74 (0.62-0.89) | 0.74 (0.62-0.87) | 0.74 (0.62-0.90) |
| Z1b | 0.74 (0.56-1.22) | 0.87 (0.70-1.08) | 1.00 (0.84-1.45) | 1.00 (0.76-1.19) |
| Z2a | 0.58 (0.43-0.85) | 0.81 (0.68-0.97) | 0.87 (0.71-1.16) | 0.82 (0.63-1.02) |
| Z2b | 0.56 (0.46-0.70) | 0.55 (0.47-0.63) | 0.55 (0.44-0.71) | 0.58 (0.47-0.68) |
| Z3 | 0.58 (0.48-0.80) | 0.73 (0.60-0.87) | 0.89 (0.74-1.04) | 0.82 (0.68-0.98) |

Note – The different Z chromosomes are divided into a and b regions according to previously inferred stepwise fusions of autosomal parts to the Z chromosomes. Confidence intervals in brackets were obtained with stratified bootstrap.

**Supplementary table 11.** Results of pairwise Wilcoxon rank sum tests of difference in gene expression levels between *unbiased* Z-linked genes and *unbiased* autosomal genes for **A**, *L. juvernica* female adult abdomen; **B**, *L. juvernica* male adult abdomen; **C**, *L. sinapis* female adult abdomen; **D**; *L. sinapis* male adult abdomen; **E**, *L. reali* female adult abdomen; **F**, *L. reali* male adult abdomen. Significant test p-values are highlighted in bold.

**A**

| group1 | group2 | n1 | n2 | statistic | p-value | p.adj | p.adj.signif |
|---|---|---|---|---|---|---|---|
| A | Z1a | 5,786 | 221 | 782622 | 1.49e-08 | **2.47e-07** | *** |
| A | Z1b | 5,786 | 107 | 280620 | 0.097 | 0.403 | |
| A | Z2a | 5,786 | 131 | 384386 | 0.78 | 1 | |
| A | Z2b | 5,786 | 155 | 592538 | 7.97e-12 | **3.97e-10** | *** |
| A | Z3 | 5,786 | 159 | 502006 | 0.049 | 0.244 | |
| Z1a | Z1b | 221 | 107 | 8016 | 2.27e-06 | **2.75e-05** | *** |
| Z1a | Z2a | 221 | 131 | 11313 | 6.12e-4 | **0.004** | ** |
| Z1a | Z2b | 221 | 155 | 19096 | 0.058 | 0.262 | |
| Z1a | Z3 | 221 | 159 | 14826 | 0.009 | 0.052 | |
| Z1b | Z2a | 107 | 131 | 7768 | 0.151 | 0.578 | |

| | | | | | | | |
|---|---|---|---|---|---|---|---|
| Z1b | Z2b | 107 | 155 | 11869 | 3,00E-09 | **7.47e-08** | *** |
| Z1b | Z3 | 107 | 159 | 10110 | 0.009 | 0.052 | |
| Z2a | Z2b | 131 | 155 | 13420 | 2.76e-06 | **2.75e-05** | *** |
| Z2a | Z3 | 131 | 159 | 11211 | 0.263 | 0.935 | |
| Z2b | Z3 | 155 | 159 | 8982 | 3.29e-05 | 2.73e-4 | *** |

**B**

| group1 | group2 | n1 | n2 | statistic | p-value | p.adj | p.adj.signif |
|---|---|---|---|---|---|---|---|
| A | Z1a | 5,789 | 224 | 773707 | 8.8e-07 | **9.7e-06** | *** |
| A | Z1b | 5,789 | 107 | 268300 | 0.018 | 0.088 | |
| A | Z2a | 5,789 | 131 | 377318 | 0.923 | 1 | |
| A | Z2b | 5,789 | 155 | 594289 | 4.93e-12 | **2.45e-10** | *** |
| A | Z3 | 5,789 | 158 | 496039 | 0.069 | 0.287 | |
| Z1a | Z1b | 224 | 107 | 7996 | 9.74e-07 | **9.7e-06** | *** |
| Z1a | Z2a | 224 | 131 | 11733 | 0.002 | **0.012** | * |
| Z1a | Z2b | 224 | 155 | 19931 | 0.014 | 0.079 | |
| Z1a | Z3 | 224 | 158 | 15443 | 0.034 | 0.154 | |
| Z1b | Z2a | 107 | 131 | 7936 | 0.079 | 0.304 | |
| Z1b | Z2b | 107 | 155 | 12251 | 5.21e-11 | **1.3e-09** | *** |
| Z1b | Z3 | 107 | 158 | 10321 | 0.002 | **0.014** | * |
| Z2a | Z2b | 131 | 155 | 13603 | 7.39e-07 | **9.7e-06** | *** |
| Z2a | Z3 | 131 | 158 | 11334 | 0.164 | 0.583 | |
| Z2b | Z3 | 155 | 158 | 8768 | 1.41e-05 | 1.17e-4 | *** |

**C**

| group1 | group2 | n1 | n2 | statistic | p-value | p.adj | p.adj.signif |
|---|---|---|---|---|---|---|---|
| A | Z1a | 5,328 | 257 | 865797 | 7.23e-13 | **3.6e-11** | *** |
| A | Z1b | 5,328 | 107 | 285115 | 0.997 | 1 | |
| A | Z2a | 5,328 | 124 | 347316 | 0.327 | 1 | |
| A | Z2b | 5,328 | 138 | 450221 | 6.41e-06 | **1.60e-04** | *** |
| A | Z3 | 5,328 | 155 | 454972 | 0.03 | 0.189 | |
| Z1a | Z1b | 257 | 107 | 9826 | 1.79e-05 | **2.97e-04** | *** |
| Z1a | Z2a | 257 | 124 | 12555 | 7.96e-4 | **0.01** | ** |
| Z1a | Z2b | 257 | 138 | 16921 | 0.453 | 1 | |
| Z1a | Z3 | 257 | 155 | 16391 | 0.003 | **0.022** | * |
| Z1b | Z2a | 107 | 124 | 7016 | 0.451 | 1 | |
| Z1b | Z2b | 107 | 138 | 9168 | 0.001 | **0.012** | * |
| Z1b | Z3 | 107 | 155 | 9192 | 0.136 | 0.677 | |
| Z2a | Z2b | 124 | 138 | 10076 | 0.013 | 0.093 | |
| Z2a | Z3 | 124 | 155 | 10101 | 0.464 | 1 | |
| Z2b | Z3 | 138 | 155 | 9279 | 0.051 | 0.279 | |

**D**

| group1 | group2 | n1 | n2 | statistic | p-value | p.adj | p.adj.signif |
|---|---|---|---|---|---|---|---|
| A | Z1a | 5,332 | 256 | 842708 | 2.1e-10 | **1.05e-08** | *** |
| A | Z1b | 5,332 | 107 | 283385 | 0.907 | 1 | |
| A | Z2a | 5,332 | 126 | 357434 | 0.218 | 0.986 | |
| A | Z2b | 5,332 | 138 | 453827 | 2.72e-06 | **6.77e-05** | *** |
| A | Z3 | 5,332 | 155 | 452482 | 0.044 | 0.241 | |
| Z1a | Z1b | 256 | 107 | 10302 | 1.97e-4 | **0.003** | ** |
| Z1a | Z2a | 256 | 126 | 13455 | 0.008 | 0.084 | |
| Z1a | Z2b | 256 | 138 | 17675 | 0.992 | 1 | |
| Z1a | Z3 | 256 | 155 | 16872 | 0.011 | 0.091 | |
| Z1b | Z2a | 107 | 126 | 7257 | 0.315 | 1 | |
| Z1b | Z2b | 107 | 138 | 9194 | 9.99e-4 | **0.012** | * |
| Z1b | Z3 | 107 | 155 | 9146 | 0.157 | 0.781 | |
| Z2a | Z2b | 126 | 138 | 10160 | 0.018 | 0.128 | |
| Z2a | Z3 | 126 | 155 | 10049 | 0.676 | 1 | |
| Z2b | Z3 | 138 | 155 | 9089 | 0.027 | 0.165 | |

**E**

| group1 | group2 | n1 | n2 | statistic | p-value | p.adj | p.adj.signif |
|---|---|---|---|---|---|---|---|
| A | Z1a | 4,563 | 195 | 533393 | 2.46e-06 | **1.22e-4** | *** |
| A | Z1b | 4,563 | 98 | 238175 | 0.268 | 1 | |
| A | Z2a | 4,563 | 112 | 273855 | 0.194 | 1 | |
| A | Z2b | 4,563 | 160 | 431868 | 8.08e-05 | **0.002** | ** |
| A | Z3 | 4,563 | 139 | 362605 | 0.004 | 0.065 | |
| Z1a | Z1b | 195 | 98 | 8322 | 0.072 | 0.714 | |
| Z1a | Z2a | 195 | 112 | 9540 | 0.065 | 0.714 | |
| Z1a | Z2b | 195 | 160 | 15394 | 0.831 | 1 | |
| Z1a | Z3 | 195 | 139 | 12660 | 0.305 | 1 | |
| Z1b | Z2a | 98 | 112 | 5569 | 0.855 | 1 | |
| Z1b | Z2b | 98 | 160 | 8702 | 0.139 | 0.988 | |
| Z1b | Z3 | 98 | 139 | 7276 | 0.372 | 1 | |
| Z2a | Z2b | 112 | 160 | 9984 | 0.109 | 0.904 | |
| Z2a | Z3 | 112 | 139 | 8286 | 0.38 | 1 | |
| Z2b | Z3 | 160 | 139 | 10502 | 0.408 | 1 | |

**F**

| group1 | group2 | n1 | n2 | statistic | p-value | p.adj | p.adj.signif |
|---|---|---|---|---|---|---|---|
| A | Z1a | 4,569 | 195 | 524751 | 2.5e-05 | **6.22e-4** | *** |
| A | Z1b | 4,569 | 98 | 234231 | 0.433 | 1 | |
| A | Z2a | 4,569 | 112 | 271155 | 0.279 | 1 | |
| A | Z2b | 4,569 | 160 | 445837 | 2.23e-06 | **1.11e-4** | *** |
| A | Z3 | 4,569 | 138 | 356929 | 0.008 | 0.134 | |
| Z1a | Z1b | 195 | 98 | 8329 | 0.073 | 0.548 | |
| Z1a | Z2a | 195 | 112 | 9596 | 0.077 | 0.548 | |
| Z1a | Z2b | 195 | 160 | 16331 | 0.448 | 1 | |
| Z1a | Z3 | 195 | 138 | 12672 | 0.366 | 1 | |
| Z1b | Z2a | 98 | 112 | 5590 | 0.817 | 1 | |
| Z1b | Z2b | 98 | 160 | 9187 | 0.021 | 0.205 | |
| Z1b | Z3 | 98 | 138 | 7298 | 0.3 | 1 | |
| Z2a | Z2b | 112 | 160 | 10446 | 0.02 | 0.205 | |
| Z2a | Z3 | 112 | 138 | 8288 | 0.325 | 1 | |
| Z2b | Z3 | 160 | 138 | 9842 | 0.106 | 0.659 | |

Correction for multiple testing was done with Benjamini-Yekutieli correction. Significant adjusted p-values are highlighted in bold and asterisks denote significance levels (* P ≤ 0.05, ** P ≤ 0.01, *** P ≤ 0.001).

**Supplementary table 12.** Female unique SNPs, intersected across all sample groups.

| Species | Scaffold | Ancestral Leptidea Linkage | SNP count | Gene count |
|---|---|---|---|---|
| *L. juvernica* | HiC_scaffold_30 | Z3 | 153 | 35 |
| | HiC_scaffold_34 | Z3 | 121 | 26 |
| *L. sinapis* | HiC_scaffold_18 | Z3 | 331 | 59 |
| | HiC_scaffold_21 | Autosome | 6 | 1 |
| | HiC_scaffold_11 | Autosome | 1 | 1 |
| *L. reali* | HiC_scaffold_2 | Z3 | 323 | 50 |
| | HiC_scaffold_24 | Autosome | 1 | 1 |
| | HiC_scaffold_25 | Autosome | 6 | 6 |
| | HiC_scaffold_11 | Autosome | 2 | 1 |
| | HiC_scaffold_3 | Autosome | 2 | 1 |

**Supplementary table 13.** Female unique SNPs, identified separately for each sample group, in each case compared against all male samples.

| Species | Stage | Tissue | Scaffold | Ancestral Leptidea Linkage | SNP count | Gene count |
|---|---|---|---|---|---|---|
| *L. juvernica* | larva | abdomen | HiC_scaffold_30 | Z3 | 285 | 44 |
| | | | HiC_scaffold_9 | Autosome | 4 | 1 |
| | | | HiC_scaffold_34 | Z3 | 285 | 35 |
| | | head | HiC_scaffold_30 | Z3 | 421 | 56 |
| | | | HiC_scaffold_7 | Autosome | 2 | 1 |
| | | | HiC_scaffold_8 | Autosome | 4 | 1 |
| | | | HiC_scaffold_9 | Autosome | 4 | 1 |
| | | | HiC_scaffold_34 | Z3 | 406 | 45 |
| | adult | abdomen | HiC_scaffold_2 | Autosome | 2 | 1 |
| | | | HiC_scaffold_30 | Z3 | 277 | 49 |
| | | | HiC_scaffold_7 | Autosome | 1 | 1 |
| | | | HiC_scaffold_8 | Autosome | 4 | 1 |
| | | | HiC_scaffold_34 | Z3 | 207 | 38 |
| | | | HiC_scaffold_36 | Autosome | 1 | 1 |
| | | | HiC_scaffold_18 | Autosome | 1 | 1 |
| | | head | HiC_scaffold_30 | Z3 | 334 | 56 |
| | | | HiC_scaffold_7 | Autosome | 1 | 1 |
| | | | HiC_scaffold_34 | Z3 | 231 | 40 |
| *L. sinapis* | larva | abdomen | HiC_scaffold_11 | Autosome | 1 | 1 |
| | | | HiC_scaffold_21 | Autosome | 6 | 1 |
| | | | HiC_scaffold_18 | Z3 | 471 | 75 |
| | | head | HiC_scaffold_11 | Autosome | 3 | 1 |
| | | | HiC_scaffold_21 | Autosome | 12 | 1 |
| | | | HiC_scaffold_18 | Z3 | 653 | 91 |
| | adult | abdomen | HiC_scaffold_2 | Autosome | 1 | 1 |
| | | | HiC_scaffold_3 | Autosome | 2 | 1 |
| | | | HiC_scaffold_11 | Autosome | 5 | 2 |
| | | | HiC_scaffold_21 | Autosome | 15 | 1 |
| | | | HiC_scaffold_7 | Autosome | 1 | 1 |
| | | | HiC_scaffold_8 | Autosome | 1 | 2 |
| | | | HiC_scaffold_18 | Z3 | 723 | 95 |
| | | head | HiC_scaffold_11 | Autosome | 1 | 1 |
| | | | HiC_scaffold_21 | Autosome | 9 | 1 |
| | | | HiC_scaffold_18 | Z3 | 729 | 98 |
| *L. reali* | larva | abdomen | HiC_scaffold_2 | Z3 | 385 | 58 |
| | | | HiC_scaffold_3 | Autosome | 2 | 1 |
| | | | HiC_scaffold_11 | Autosome | 2 | 1 |
| | | | HiC_scaffold_24 | Autosome | 7 | 1 |
| | | | HiC_scaffold_25 | Autosome | 7 | 7 |
| | | head | HiC_scaffold_2 | Z3 | 640 | 84 |
| | | | HiC_scaffold_3 | Autosome | 2 | 1 |
| | | | HiC_scaffold_11 | Autosome | 10 | 1 |
| | | | HiC_scaffold_24 | Autosome | 13 | 1 |
| | | | HiC_scaffold_15 | Autosome | 2 | 1 |
| | | | HiC_scaffold_25 | Autosome | 20 | 13 |
| | adult | abdomen | HiC_scaffold_1 | Autosome | 2 | 1 |
| | | | HiC_scaffold_2 | Z3 | 613 | 74 |
| | | | HiC_scaffold_3 | Autosome | 4 | 2 |

| | | | | | |
|---|---|---|---|---|---|
| | | HiC_scaffold_11 | Autosome | 7 | 1 |
| | | HiC_scaffold_6 | Autosome | 5 | 2 |
| | | HiC_scaffold_24 | Autosome | 13 | 1 |
| | | HiC_scaffold_25 | Autosome | 22 | 17 |
| | head | HiC_scaffold_2 | Z3 | 782 | 98 |
| | | HiC_scaffold_3 | Autosome | 2 | 1 |
| | | HiC_scaffold_11 | Autosome | 10 | 1 |
| | | HiC_scaffold_24 | Autosome | 16 | 2 |
| | | HiC_scaffold_15 | Autosome | 2 | 1 |
| | | HiC_scaffold_25 | Autosome | 27 | 19 |

**Supplementary table 14.** Test results for Pearsons correlation between M/F gene expression log2 fold change and the expression ratio of the reference Z allele. Significant test p-values are highlighted in bold. P-values were corrected using the Benjamini-Hochberg method.

| species | sex | stage | tissue | statistic | correlation | p-value | p.adj |
|---|---|---|---|---|---|---|---|
| *L. juvernica* | female | adult | abdomen | 1.8382 | 0.197 | 0.070 | 0.152 |
| | | | head | 8.9474 | 0.680 | 3.448e-14 | **4.14e-13** |
| | | larva | abdomen | 7.3228 | 0.643 | 2.154e-10 | **1.72e-09** |
| | | | head | 5.7188 | 0.500 | 1.163e-07 | **4.65e-07** |
| | male | adult | abdomen | -0.70994 | -0.080 | 0.480 | 0.823 |
| | | | head | 0.049778 | 0.005 | 0.960 | 0.961 |
| | | larva | abdomen | -3.2099 | -0.354 | 0.002 | **0.005** |
| | | | head | -0.42293 | -0.044 | 0.673 | 0.951 |
| *L. sinapis* | female | adult | abdomen | 1.3596 | 0.140 | 0.177 | 0.327 |
| | | | head | 6.9772 | 0.584 | 4.18e-10 | **2.51e-09** |
| | | larva | abdomen | 3.7691 | 0.406 | 3.327e-4 | **8.87e-04** |
| | | | head | 4.81 | 0.456 | 6.197e-06 | **2.12e-05** |
| | male | adult | abdomen | 0.34301 | 0.036 | 0.732 | 0.961 |
| | | | head | 0.048762 | 0.005 | 0.961 | 0.961 |
| | | larva | abdomen | 0.46794 | 0.056 | 0.641 | 0.951 |
| | | | head | -0.5617 | -0.060 | 0.576 | 0.921 |
| *L. reali* | female | adult | abdomen | 1.6539 | 0.191 | 0.103 | 0.205 |
| | | | head | 9.803 | 0.707 | 3.942e-16 | **9.46e-15** |
| | | larva | abdomen | 4.2141 | 0.491 | 9.216e-05 | **2.77e-04** |
| | | | head | 6.442 | 0.580 | 7.568e-09 | **3.63e-08** |
| | male | adult | abdomen | -0.23554 | -0.028 | 0.815 | 0.961 |
| | | | head | -0.062177 | -0.006 | 0.951 | 0.961 |
| | | larva | abdomen | -0.17642 | -0.024 | 0.861 | 0.961 |
| | | | head | -0.1215 | -0.014 | 0.903 | 0.961 |

**Supplementary table 15.** Test results for Pearson correlation between gene expression levels (TPM) and the expression ratio of the reference Z allele. Significant test p-values are highlighted in bold. P-values were corrected using the Benjamini-Hochberg method.

| species | sex | stage | tissue | statistic | correlation | p-value | p.adj |
|---|---|---|---|---|---|---|---|
| *L. juvernica* | female | adult | abdomen | -0.96992 | -0.105 | 0.335 | 0.536 |
| | | | head | 0.091622 | 0.010 | 0.927 | 0.979 |
| | | larva | abdomen | 0.98141 | 0.112 | 0.330 | 0.536 |
| | | | head | 1.0102 | 0.102 | 0.315 | 0.536 |
| | male | adult | abdomen | 3.2292 | 0.341 | 0.002 | **0.034** |
| | | | head | 2.1577 | 0.225 | 0.034 | 0.202 |
| | | larva | abdomen | 1.141 | 0.133 | 0.258 | 0.515 |

| | | | | | | | |
|---|---|---|---|---|---|---|---|
| | | | head | 2.2548 | 0.228 | 0.026 | 0.202 |
| *L. sinapis* | female | adult | abdomen | -1.2185 | -0.126 | 0.226 | 0.493 |
| | | | head | -0.2314 | -0.024 | 0.818 | 0.934 |
| | | larva | abdomen | 1.3656 | 0.159 | 0.176 | 0.493 |
| | | | head | 0.067971 | 0.007 | 0.946 | 0.979 |
| | male | adult | abdomen | 1.6985 | 0.176 | 0.093 | 0.371 |
| | | | head | 1.2743 | 0.131 | 0.206 | 0.493 |
| | | larva | abdomen | -0.026598 | -0.003 | 0.979 | 0.979 |
| | | | head | 1.7386 | 0.183 | 0.086 | 0.371 |
| *L. reali* | female | adult | abdomen | 0.41018 | 0.048 | 0.683 | 0.819 |
| | | | head | -0.85314 | -0.087 | 0.396 | 0.594 |
| | | larva | abdomen | 0.42727 | 0.057 | 0.671 | 0.819 |
| | | | head | -0.60859 | -0.067 | 0.545 | 0.726 |
| | male | adult | abdomen | 1.2269 | 0.145 | 0.224 | 0.493 |
| | | | head | 1.5571 | 0.159 | 0.123 | 0.421 |
| | | larva | abdomen | -0.77744 | -0.105 | 0.440 | 0.622 |
| | | | head | 3.0837 | 0.326 | 0.003 | **0.034** |

**Supplementary table 16.** Results of Wilcoxon rank sum test of significant difference in gene expression levels (TPM) in males between the gene set that showed monoallelic expression in females, and male autosomal expression levels. 'A' and 'Z/Z' shows number of genes in each test, 'Z/Z' in this case representing the genes with 'Z/0' expression in females. P-values were corrected using the Benjamini-Yekutieli method. Significant test p-values are highlighted in bold.

| species | stage | tissue | A | Z/Z | statistic | p-value | p.adj |
|---|---|---|---|---|---|---|---|
| *L. juvernica* | adult | abdomen | 10308 | 223 | 1,329,906 | 5.820e-05 | **3.61e-04** |
| | | head | 10149 | 203 | 1,277,993 | 4.122e-09 | **5.12e-08** |
| | larva | Abdomen | 9516 | 215 | 1,153,203 | 1.388e-3 | **0.005** |
| | | head | 9384 | 191 | 1,035,304 | 2.343e-4 | **9.69e-04** |
| *L. sinapis* | adult | abdomen | 10003 | 198 | 1,164,594 | 2.161e-05 | **1.61e-04** |
| | | head | 9791 | 187 | 1,163,087 | 2.207e-10 | **4.11e-09** |
| | larva | abdomen | 9169 | 189 | 947,194 | 0.028 | 0.087 |
| | | head | 9057 | 168 | 858,563 | 4.254e-3 | **0.014** |
| *L. reali* | adult | abdomen | 9618 | 216 | 1,200,772 | 8.613e-05 | **4.01e-04** |
| | | head | 9463 | 182 | 1,126,747 | 9.424e-13 | **3.51e-11** |
| | larva | abdomen | 8977 | 209 | 1,087,315 | 8.244e-5 | **4.01e-04** |
| | | head | 8842 | 176 | 976,133 | 7.013e-09 | **6.53e-08** |

**Supplementary table 17.** Results of Wilcoxon rank sum test of significant difference in gene expression levels (TPM) between genes with monoallelic and bi-allelic expression in females. 'Z/0' and 'Z/W' shows number of genes in each test. P-values were corrected using the Benjamini-Hochberg method. Significant test p-values are highlighted in bold.

| species | stage | tissue | Z/0 | Z/W | statistic | p-value | p.adj |
|---|---|---|---|---|---|---|---|
| *L. juvernica* | adult | abdomen | 224 | 86 | 2,747 | 1.97e-22 | **1.182e-21** |
| | | head | 212 | 95 | 3,602 | 2.35e-19 | **4.7e-19** |
| | larva | abdomen | 208 | 78 | 3,234 | 4.87e-15 | **7.305e-15** |
| | | head | 192 | 100 | 4,633 | 4.06e-13 | **4.872e-13** |
| *L. sinapis* | adult | abdomen | 194 | 94 | 2,985 | 2.17e-20 | **5.208e-20** |
| | | head | 190 | 96 | 2,522 | 1.71e-23 | **2.052e-22** |
| | larva | abdomen | 184 | 74 | 3,091 | 7.1e-12 | **7.745e-12** |
| | | head | 168 | 90 | 3,972 | 3.39e-10 | **3.39e-10** |
| *L. reali* | adult | abdomen | 214 | 74 | 2,051 | 2.11e-21 | **6.33e-21** |
| | | head | 185 | 98 | 2,763 | 6.58e-22 | **2.632e-21** |

| | | | | | | | |
|---|---|---|---|---|---|---|---|
| | larva | abdomen | 201 | 58 | 1,918 | 7.21e-15 | **9.613e-15** |
| | | head | 180 | 84 | 2,460 | 1.09e-18 | **1.869e-18** |

**Supplementary table 18.** Significantly overrepresented GO terms for Z chromosome regions with male downregulation. A, *L. juvernica*; B, *L. sinapis*; C, *L. reali.*

**A**

| GO.ID | Term | Annotated | Significant | Expected | P-value |
|---|---|---|---|---|---|
| GO:0005975 | carbohydrate metabolic process | 25 | 22 | 14.62 | 0.0072 |
| GO:0007186 | G protein-coupled receptor signaling pat... | 13 | 12 | 7.6 | 0.0351 |
| GO:0007018 | microtubule-based movement | 7 | 7 | 4.09 | 0.0387 |
| GO:0016020 | membrane | 125 | 78 | 68.46 | 0.041 |
| GO:0005874 | microtubule | 7 | 7 | 3.83 | 0.014 |
| GO:0003700 | DNA-binding transcription factor activit... | 19 | 17 | 12.1 | 0.01199 |
| GO:0003676 | nucleic acid binding | 196 | 143 | 124.84 | 0.00092 |
| GO:0016758 | hexosyltransferase activity | 25 | 18 | 15.92 | 0.02575 |
| GO:0004930 | G protein-coupled receptor activity | 11 | 11 | 7.01 | 0.01660 |
| GO:0004553 | hydrolase activity, hydrolyzing O-glycos... | 17 | 15 | 10.83 | 0.02601 |

**B**

| GO.ID | Term | Annotated | Significant | Expected | P-value |
|---|---|---|---|---|---|
| GO:0005975 | carbohydrate metabolic process | 24 | 20 | 13.67 | 0.034 |
| GO:0006355 | regulation of DNA-templated transcriptio... | 39 | 28 | 22.22 | 0.017 |
| GO:0016020 | membrane | 135 | 89 | 74.39 | 0.0037 |
| GO:0003700 | DNA-binding transcription factor activit... | 17 | 15 | 10.57 | 0.01819 |
| GO:0003676 | nucleic acid binding | 188 | 134 | 116.9 | 0.00093 |
| GO:0016758 | hexosyltransferase activity | 20 | 14 | 12.44 | 0.04148 |

**C**

| GO.ID | Term | Annotated | Significant | Expected | P-value |
|---|---|---|---|---|---|
| GO:0005975 | carbohydrate metabolic process | 26 | 23 | 15.6 | 0.018 |
| GO:0005874 | microtubule | 6 | 6 | 3.3 | 0.027 |
| GO:0003700 | DNA-binding transcription factor activit... | 19 | 18 | 11.95 | 0.0025 |
| GO:0004930 | G protein-coupled receptor activity | 9 | 9 | 5.66 | 0.0239 |

**Supplementary table 19.** Significantly overrepresented GO terms for Z chromosome regions with female upregulation. A, *L. juvernica*; B, *L. sinapis*; C, *L. reali.*

**A**

| GO.ID | Term | Annotated | Significant | Expected | P-value |
|---|---|---|---|---|---|
| GO:0008272 | sulfate transport | 6 | 6 | 1.93 | 0.001 |
| GO:0018095 | protein polyglutamylation | 3 | 3 | 0.97 | 0.033 |
| GO:0019991 | septate junction assembly | 3 | 3 | 0.97 | 0.033 |
| GO:0006629 | lipid metabolic process | 11 | 9 | 3.55 | 0.035 |
| GO:0005615 | extracellular space | 5 | 4 | 1.68 | 0.045 |
| GO:0008271 | secondary active sulfate transmembrane t... | 6 | 6 | 1.72 | 0.00054 |
| GO:0008146 | sulfotransferase activity | 3 | 3 | 0.86 | 0.02354 |
| GO:0004402 | histone acetyltransferase activity | 3 | 3 | 0.86 | 0.02354 |
| GO:0005242 | inward rectifier potassium channel activ... | 3 | 3 | 0.86 | 0.02354 |
| GO:0008417 | fucosyltransferase activity | 5 | 4 | 1.44 | 0.02591 |
| GO:0004252 | serine-type endopeptidase activity | 21 | 10 | 6.04 | 0.04959 |

**B**

| GO.ID | Term | Annotated | Significant | Expected | P-value |
|---|---|---|---|---|---|
| GO:0008272 | sulfate transport | 7 | 7 | 2.28 | 0.00036 |
| GO:0018095 | protein polyglutamylation | 4 | 4 | 1.31 | 0.01106 |
| GO:0019991 | septate junction assembly | 3 | 3 | 0.98 | 0.03433 |
| GO:0005615 | extracellular space | 5 | 4 | 1.7 | 0.047 |
| GO:0005576 | extracellular region | 15 | 11 | 5.1 | 0.018 |
| GO:0008271 | secondary active sulfate transmembrane t... | 7 | 7 | 1.96 | 0.00013 |
| GO:0008146 | sulfotransferase activity | 3 | 3 | 0.84 | 0.02184 |
| GO:0004402 | histone acetyltransferase activity | 3 | 3 | 0.84 | 0.02184 |
| GO:0005242 | inward rectifier potassium channel activ... | 3 | 3 | 0.84 | 0.02184 |
| GO:0008417 | fucosyltransferase activity | 5 | 4 | 1.4 | 0.02362 |
| GO:0004252 | serine-type endopeptidase activity | 20 | 11 | 5.61 | 0.00927 |

| | | | | | |
|---|---|---|---|---|---|
| GO:0003735 | structural constituent of ribosome | 10 | 6 | 2.8 | 0.03355 |
| GO:0042302 | structural constituent of cuticle | 7 | 7 | 1.96 | 0.00013 |

**C**

| GO.ID | Term | Annotated | Significant | Expected | P-value |
|---|---|---|---|---|---|
| GO:0008272 | sulfate transport | 6 | 6 | 1.82 | 0.00074 |
| GO:0018095 | protein polyglutamylation | 4 | 4 | 1.22 | 0.00832 |
| GO:0019991 | septate junction assembly | 4 | 4 | 1.22 | 0.00832 |
| GO:0006629 | lipid metabolic process | 13 | 9 | 3.95 | 0.02432 |
| GO:0070588 | calcium ion transmembrane transport | 3 | 3 | 0.91 | 0.02774 |
| GO:0006486 | protein glycosylation | 10 | 6 | 3.04 | 0.04866 |
| GO:0005615 | extracellular space | 4 | 4 | 1.34 | 0.012 |
| GO:0008271 | secondary active sulfate transmembrane t... | 6 | 6 | 1.73 | 0.00055 |
| GO:0008146 | sulfotransferase activity | 3 | 3 | 0.87 | 0.02374 |
| GO:0004402 | histone acetyltransferase activity | 3 | 3 | 0.87 | 0.02374 |
| GO:0003735 | structural constituent of ribosome | 8 | 5 | 2.31 | 0.04846 |

**Supplementary table 20.** Significantly overrepresented GO terms for genes with expression of female unique SNPs on Z3, indicative of W gametolog expression. A, *L. sinapis*; B, *L. reali.*

**A**

| GO.ID | Term | Annotated | Significant | Expected | P-value |
|---|---|---|---|---|---|
| GO:0003676 | nucleic acid binding | 37 | 17 | 13.26 | 0.034 |

**B**

| GO.ID | Term | Annotated | Significant | Expected | P-value |
|---|---|---|---|---|---|
| GO:0003676 | nucleic acid binding | 36 | 16 | 11.78 | 0.0106 |
| GO:0008270 | zinc ion binding | 7 | 6 | 2.29 | 0.0052 |
